# Profilin promotes lamellipodium protrusion by tuning the antagonistic activities of CP and VASP

**DOI:** 10.1101/2025.03.26.645585

**Authors:** Yubo Tang, Matthias Schaks, Magdalena Mietkowska, Jonas Scholz, Ruth Benavente-Naranjo, Sarah Körber, Zhilun Li, Christopher Lambert, Theresia E.B. Stradal, Roger Karlsson, Peter Bieling, Jan Faix, Martin Falcke, Klemens Rottner

## Abstract

Cell migration on rigid surfaces employs flat protrusions termed lamellipodia, which constitute the prime model system for branched actin filament networks that generate pushing forces for membrane movement. Branched actin filaments are nucleated by the Arp2/3 complex and play vital roles in various cell biological processes, such as organelle trafficking, phagocytosis and autophagy. Here we utilize genome editing to explore the functional connections between the actin monomer-binding protein profilin (Pfn), the filament nucleating Arp2/3 complex, its co-factor heterodimeric capping protein (CP) and the Ena/VASP family of actin filament polymerases in lamellipodial actin assembly. Individual and combinations of knockouts show that Pfn counters Ena/VASP but promotes Arp2/3 complex activity, while Ena/VASP and CP mutually antagonize each other. Notably, while Pfn is important for Arp2/3 complex activity irrespective of Ena/VASP, sensitivity of CP to Pfn removal is lost in the absence of Ena/VASP. Our findings establish Pfn as master regulator of Arp2/3 complex-dependent actin network formation, which differentially regulates VASP and its antagonizer CP. Finally, mathematical modeling of our data suggest that Ena/VASP and CP compete for binding at the lamellipodial edge, likely contributing to their functional antagonism at this subcellular site. This work provides critical insights into the molecular logic of branched actin network assembly in membrane protrusion and cellular force generation.

## Introduction

Actin filament assembly is essential for numerous cellular processes such as shape establishment and change, division, organelle trafficking as well as cell-cell and cell-environment interactions. Actin filaments are continuously assembled from ATP-bound monomers, which are tightly regulated by conserved monomer-binding proteins. However, most research in past decades has focused on factors promoting filament nucleation (Cao & Way, 2024; Gautreau *et al*, 2022; Rottner *et al*, 2017). Cells construct diverse actin structures depending on type, physiological state as well as tissue context. Actin filaments in these structures constantly polymerize and depolymerize, but the rate of turnover differs depending not only on network type, but also cell type and function. For example, actin networks at the leading edge of moving cells are more dynamic than those found in the interior of static cells, which feature large, rigid structures such as stress fibers anchored at focal adhesions (Dimchev & Rottner, 2018; Saito *et al*, 2023).

One of the most dynamic actin structures is the lamellipodium, crucial for cell migration and axonal growth (Leite *et al*, 2021; Rottner & Schaks, 2019; Sadhu *et al*, 2023). It serves as key model system for studying rapid actin remodeling, due to its consistent features, such as a network density gradient from front to back and a uniform criss-cross arrangement of branched actin filaments. Arp2/3 complex activation at the lamellipodium front generates a dense array of branched filaments that create pushing forces, which drive cell protrusion. The WAVE Regulatory Complex (WRC), activated by the Rho-family GTPase Rac (1, 2 or 3), is a major activator of Arp2/3 complex, leading to branching at the lamellipodium front (Alekhina *et al*, 2017; Bieling & Rottner, 2023). Despite the central role of the WH2-, central and acidic motifs (WCA) in autoinhibitory interactions of the WRC and the activation of the Arp2/3 complex (Rottner *et al*, 2021), recent data suggest that the Arp2/3-activating activity of the WCA domain can be replaced by alternative pathways (Buracco *et al*, 2024). Aside from continuous branching at the lamellipodium front, the interruption of which leads to lamellipodium collapse (Koestler *et al*, 2013), a limited number of additional activities are needed in order to maintain a balance between actin filament assembly and disassembly within the lamellipodium. These include actin filament polymerases such as of the Ena/VASP or formin classes of proteins (Faix & Rottner, 2022; Zimmermann & Kovar, 2019), filament capping proteins such as CP (Edwards *et al*, 2014) and actin disassembly factors such as cofilin/actin depolymerizing factor (ADF) members (Goode *et al*, 2023). The latter are essential for actin network turnover (Kanellos & Frame, 2016; Loisel *et al*, 1999), whereas CP limits filament elongation and thereby length and also indirectly promotes Arp2/3-dependent branching (Funk *et al*, 2021; Hein *et al*, 2023).

Profilin (Pfn), an actin monomer-binding protein, plays a critical, yet underexplored, role in this process. Of the four Pfn genes present in the mammalian genome, Pfn1 is most abundant and ubiquitous, whereas -2 (specifically -2a) is enriched in the nervous system and Pfn3 and -4 are-testis-specific (Karlsson & Draber, 2021). Pfn inhibits spontaneous, uncontrolled actin filament nucleation by forming a stoichiometric complex with actin monomers and promotes nucleotide exchange, thereby generating the main source of polymerizable actin available for elongation of the fast-growing -so-called barbed ends of actin filaments. It also interacts with formins to accelerate actin elongation (Kovar *et al*, 2006; Oosterheert *et al*, 2024). Though less dependent on Pfn than formins *in vitro* (Breitsprecher *et al*, 2008), Ena/VASP proteins also interact with profilin:actin, but their relationship with Pfn *in vivo* remains unclear (Faix & Rottner, 2022).

Conflicting studies have suggested that Pfn either promotes or inhibits Arp2/3 complex activity. Seminal work in fission yeast showed that Pfn favors the assembly of formin-over Arp2/3-dependent networks (Suarez *et al*, 2015). Work in mammalian cells described that Pfn overexpression inhibits lamellipodia by impeding Arp2/3 complex activity and localization at the leading edge (Rotty *et al*, 2015). Both studies thus suggested that Pfn funnels polymerizable actin into formin- or Ena/VASP-dependent structures at the expense of Arp2/3 complex-generated networks (Suarez & Kovar, 2016). However, a more recent study genetically disrupting Pfn1 expression in the neuronal CAD cell line came to opposite conclusions concerning the relation between Pfn and Arp2/3 complex functions (Skruber *et al*, 2020).

To resolve these conflicts, we investigate the impact of Pfn disruption on lamellipodia formation in B16-F1 melanoma cells, a well-established model for studying actin protrusions (Ballestrem *et al*, 1998; Mejillano *et al*, 2004; Schaks *et al*, 2018). We examine Pfn’s effects on Arp2/3 complex activity, Ena/VASP distribution, and the interaction between Pfn, Ena/VASP, and CP within lamellipodia. Mathematical modeling of Pfn-deficient and combined knockouts suggests that VASP and CP compete for binding sites at the lamellipodium front, contributing to their functional antagonism in actin network regulation.

## Results

### Profilin suppression has differential consequences on formation of lamellipodia *versus* **filopodia and formin function**

Pfn1 has previously been implicated in the formation of both lamellipodia and filopodia, for instance in the brain-tumor-derived CAD (Cath.-a-differentiated) cell line, in which the protein appeared roughly 10-fold in expression as compared to Pfn2 (Skruber *et al*., 2020). Surprisingly, Pfn1 knockout in this line was reported to be accompanied by complete removal of Arp2/3 complex from the cell periphery, and hence of canonical lamellipodia, and a clear reduction of what was termed linear actin filament arrays and supposed to correspond to microspike- or filopodia-like bundles. The described elimination of Arp2/3 complex incorporation was unexpected considering the previously proposed concept of antagonistic functions for Pfn *versus* Arp2/3 complex (Rotty *et al*., 2015; Suarez *et al*., 2015). To clarify the potential relevance of Pfn1 and overall profilin activity for lamellipodia *versus* filopodia formation or cell viability, we first disrupted the *Pfn1* gene in B16-F1 melanoma cells. Upon CRISPR/Cas9-mediated Pfn1 removal, three independently generated clones (termed Pfn1-KO #6, #10, #20) were identified by both Western blotting and sequencing of the targeted locus (Fig. 1A, B). In order to assess the potential contribution of Pfn2 to the remaining actin remodeling occurring in Pfn1-deficient cells, we first quantified the abundance of Pfn2 in these cell lines (Fig. 1C-E). To our surprise, Pfn2 expression was not upregulated, but instead even lower than WT in Pfn1-deficient cells, and expressed in a concentration range between roughly 7 and 22nM, depending on cell line. Assuming a Pfn1 concentration of 66µM, which was previously measured for the related B16-F10 cell line (Funk *et al*, 2019), this translates into expression of Pfn2 in minute amounts as compared to Pfn1, i.e. 4 to 5 orders of magnitude lower than Pfn1 in WT cells. This is consistent with the virtual elimination of mDia2 formin-dependent filopodia formation in spite of maintained subcellular positioning of the formin at the cell periphery (Supplementary Video 1). A similar abrogation was observed for filopodia induced by active variants of FMNL3 (unpublished data) and massive suppression at least of spontaneous formation of these structures in distinct culture conditions (Extended data Fig. 1). Specifically, whereas the sole frequency of lamellipodia formation was virtually identical in all Pfn1-KO *versus* WT cells (Extended Fig. 1A, B), filopodia were strongly abrogated even if induced by a combination of substratum and serum deprivation (Extended Fig. 1C, D). It is tempting to speculate that our developed lines thus constitute cells with dramatically reduced acceleration of formin-dependent actin filament elongation (Kovar *et al*., 2006), close in activity in this respect to pan-formin knockouts.

**Figure 1.**
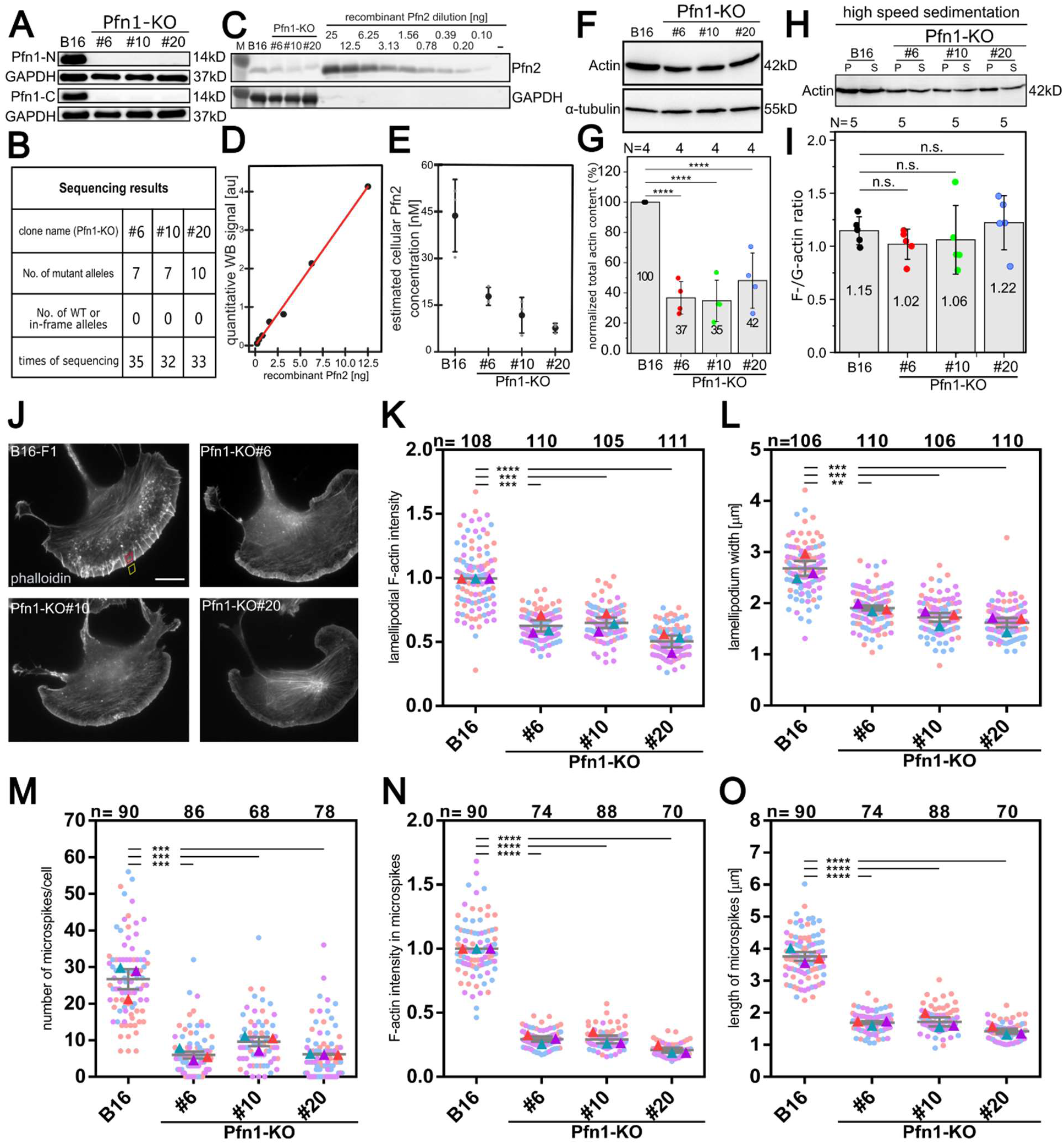
Generation and initial characterization of profilin 1 knockout (Pfn1-KO) cell lines. **(A)** Western blotting of B16-F1 cell lines before (B16) and after clonal isolation of cells treated with a Pfn1-specific guide construct (see Methods). Following initial screening, clones #6, #10 and #20 were selected for further analysis, and confirmed for the virtual absence of Pfn-1 using two independent antibodies specific for the N- and C-terminus, as indicated. GAPDH was used as loading control in each case. (**B**) Summary of the sequencing results of the genomic locus encoding the Pfn1 gene in each cell line, as indicated. No WT or in-frame deletion was detected in at least 32 independent sequencing reactions per cell line. (**C**) Representative western blot of whole-cell extracts of Pfn1-KO clones and B16-F1 cells next to a serial dilution of recombinant human profilin-2. GAPDH was used to normalize for unequal sample loading. (**D**) Example standard curve that relates the immunoblot signal to the known quantity of recombinant profilin 2 (Pfn2). The line is a linear fit to the data. (**E**) Calculated cellular concentrations of Pfn2 for WT (B16) and Pfn1-KO clones. Data from three independent experiments (gray, small symbols) were averaged to obtain mean (large, black symbols) concentrations. Error bars are standard deviations of means (SD). (**F**) Comparison of actin and tubulin expression in WT *versus* Pfn1-KO cell lines, as indicated. (**G**) Actin expression as normalized to α-tubulin and 100% in WT B16-F1 cells. Note the significant reduction of actin expression (to app. 40% on average) as compared to B16-F1 control. Data are arithmetic means and standard deviations (error bars) from four independent experiments (N). One-way ANOVA and Tukey Multiple Comparison test were used to reveal statistically significant differences between datasets. ****p≤0.001. (**H**) Representative Western blot using actin antibodies of supernatant (S) and pellet (P) fractions of respective cell lysates upon high speed sedimentation. (**I**) Quantification of the Western blotting data in H, displayed as F- over G-actin ratio (P/S-fractions) for each cell line, as indicated. Data are arithmetic means and standard deviations (error bars) from five independent experiments (N). One-way ANOVA and Tukey Multiple Comparison test were used to reveal statistically significant differences between datasets. n.s., not significant. (**J**) Representative wildtype and Pfn1-deficient cells (derived from clones as indicated) growing on laminin and fixed and stained for the actin cytoskeleton using fluorescently-coupled phalloidin. The rhomb -shaped area in red marks an example region used to measure the lamellipodial actin network intensity as shown in K. Areas lacking microspike bundles were measured, as these were measured and analyzed separately. For obtaining the data displayed in A, each area was subtracted by a close-by area outside the cell (yellow rhomb) representing extracellular background fluorescence. Scale bar: 10 µm. (**K**) Quantitation of lamellipodial actin filament (F-actin) intensities in cell lines as indicated and measured as illustrated in subpanel J. The arithmetic means of fluorescence background-subtracted B16-F1 wildtype (B16) data were normalized to 1 for each of three independent experiments, and shown as so called superplots (Lord *et al*, 2020), displaying measurements from individual cells (see Methods) and their arithmetic means as color-coded dots and triangles, respectively. Grey lines and error bars display means and standard errors of means (SEMs), n = numbers of measured cells. (**L**) Quantitation of lamellipodial widths, i.e. average orthogonal distance measured from the protruding edge toward the lamellipodial rear in the cell interior. Data assessed in principle and displayed as superplots as described in K, with the exception that absolute numbers (in µm) are shown. (**M**) Quantitation of microspike numbers embedded into lamellipodia per cell, and displayed as absolute numbers as described in L. (**N**) Quantitation of F-actin intensities throughout microspikes, with data assessed and displayed analogous to K. (**O**) Microspike lengths quantified, assessed and displayed as described for L. For all data in subpanels K-O, datasets from each individual KO-line were statistically compared to WT using two sample two-sided T-tests; **p≤0.01; ***p≤0.005; ****p≤0.001.

### Pfn elimination is cellular lethal and its suppression reduces actin expression without impacting the F-/G-actin ratio

We also sought to completely eliminate profilin activity, either by targeting Pfn2 in Pfn1-KO #20 or by disrupting Pfn1 in a stable Pfn2-KO cell line (clone #41). However, despite achieving near-complete, transient elimination of expression of the remaining profilin in each case, we were unable to obtain any stably growing clones with both genes disrupted (Extended Fig. 2A-E). Likewise, *Dictyostelium* amoeba, which harbor three functional profilins (Arasada *et al*, 2007), can be successfully and stably deprived of expression of profilins 1 and -2 (Pfn I and -II) (Haugwitz *et al*, 1994), but not, in addition, of Pfn III (Extended Fig. 2F, G), although the latter could again be efficiently deleted individually (Extended Fig. 2G, left panel and Arasada *et al*., 2007). We thus conclude that the complete absence of Pfn activity in eukaryotes is cellular lethal, likely due to crucial functions previously established for profilins, e.g. in genome stability and cytokinesis (Bottcher *et al*, 2009; Scotto di Carlo *et al*, 2023; Zhu *et al*, 2022). This would be consistent with the observation of gradually increased numbers of nuclei in our Pfn1-KO cell lines, which inversely correlated with the amount of measured Pfn2 still expressed in these lines (Fig. 1E, Extended Fig. 3).

Next, we assessed the expression of actin in our Pfn1-deficient cell lines and found it was significantly reduced to app. 40% of the level seen in WT cells (Fig. 1F, G). This reduction of actin expression did not, however, change the so called F-/G-actin ratio, i.e. the relative extent of incorporation of the remaining actin monomers into filaments (Fig. 1H, I). Clearly, this suggests that in spite of the reduction of formin-dependent actin filament elongation, the virtual lack of profilin does not, surprisingly, reduce the relative extent of cellular actin filament assembly. Although this contrasts with observations from neuronal CAD cells (Skruber *et al*., 2020), these data are consistent with the principal capability of migration (Supplementary Videos 2 and 3 for low and high magnifications, respectively) and proliferation as well as and the presence of major actin filament structures such as lamellipodia and stress fibers in our Pfn1-deficicent cell lines (Extended Fig. 1).

### Pfn loss of function causes lamellipodia to exhibit morphological and kinetic alterations and reduces protrusion and migration

To explore the precise relevance of Pfn as potential antagonist and promoter of Arp2/3-*versus* formin-induced actin assembly (Suarez & Kovar, 2016), respectively, we initially studied the precise morphological and kinetic features of lamellipodia constituting the best-characterized Arp2/3 complex-dependent structure to date (Bieling & Rottner, 2023). First, considering the potential relevance of Pfn activity and the proposal of its requirement for both Arp2/3- and Ena/VASP family member-mediated actin assembly at lamellipodial leading edges (Skruber *et al*., 2020), the protrusion of these structures and their dynamics in all three Pfn1-KO lines, as assessed by phase contrast and fluorescence video microscopy (of EGFP-tagged actin) appeared astonishingly normal (Supplementary Video 4). Phalloidin stainings of the actin cytoskeleton established lamellipodial actin networks to look at best moderately changed, except for the highly evident reduction of microspike bundles embedded into these networks (Fig. 1J). However, quantitations revealed a clear reduction of lamellipodial F-actin mass and width (Fig. 1K, L), and confirmed a dramatic suppression of microspike numbers in Pfn1-KO cells (Fig. 1M). The remaining microspikes were drastically reduced in both intensity (Fig. 1N) and length (Fig. 1O), consistent with the overall reduction of lamellipodial width (Fig. 1K). These data confirmed partially differential effects on the actin cytoskeleton already described above, with a more severe impact on protrusive bundles, such as filopodia and microspikes as compared to networks like in canonical lamellipodia. However, described morphological changes of lamellipodia also coincided with their reduced rates of protrusion (Extended Fig. 4A, B) and of lamellipodial network assembly (Extended Fig. 4C, D, Supplementary Video 5) as well as a modest abrogation of cell migration (Extended Fig. 4E, 5, Supplementary Video 2).

### Pfn loss of function reduces, but does not eliminate Arp2/3 complex activity at the cell periphery

Next, we assessed how the virtual absence of Pfn activity concomitant with the reduction of network polymerization rates in the lamellipodium impacted the distribution of its key actin assembly machineries. In a cellular cytoplasm with distinct actin filament structures, Pfn was suggested as decisive factor in funneling actin monomers into formin- over Arp2/3 complex-mediated actin assembly (Kadzik *et al*, 2020), suggesting remaining actin monomers in the absence of Pfn1 to preferentially power Arp2/3 complex-dependent actin polymerization. Contrary, however, to this proposed functional antagonism between profilin and Arp2/3 complex activities and in spite of the concomitant reduction of formin activity (see above), we found Arp2/3 complex incorporation into the lamellipodia of Pfn1-deficient cells to be significantly reduced (Fig. 2A, B), i.e. less than half on average as compared to WT cells and more severely indeed than the reduction of lamellipodial F-actin content (Fig. 1K). Numbers similar to Arp2/3 complex were obtained for its agonist CP (Fig. 2A, C), in spite of comprehensive *in vitro* data suggesting the opposite, i.e. a competition between Pfn and barbed end binders such CP (Pernier *et al*, 2016).

**Figure 2.**
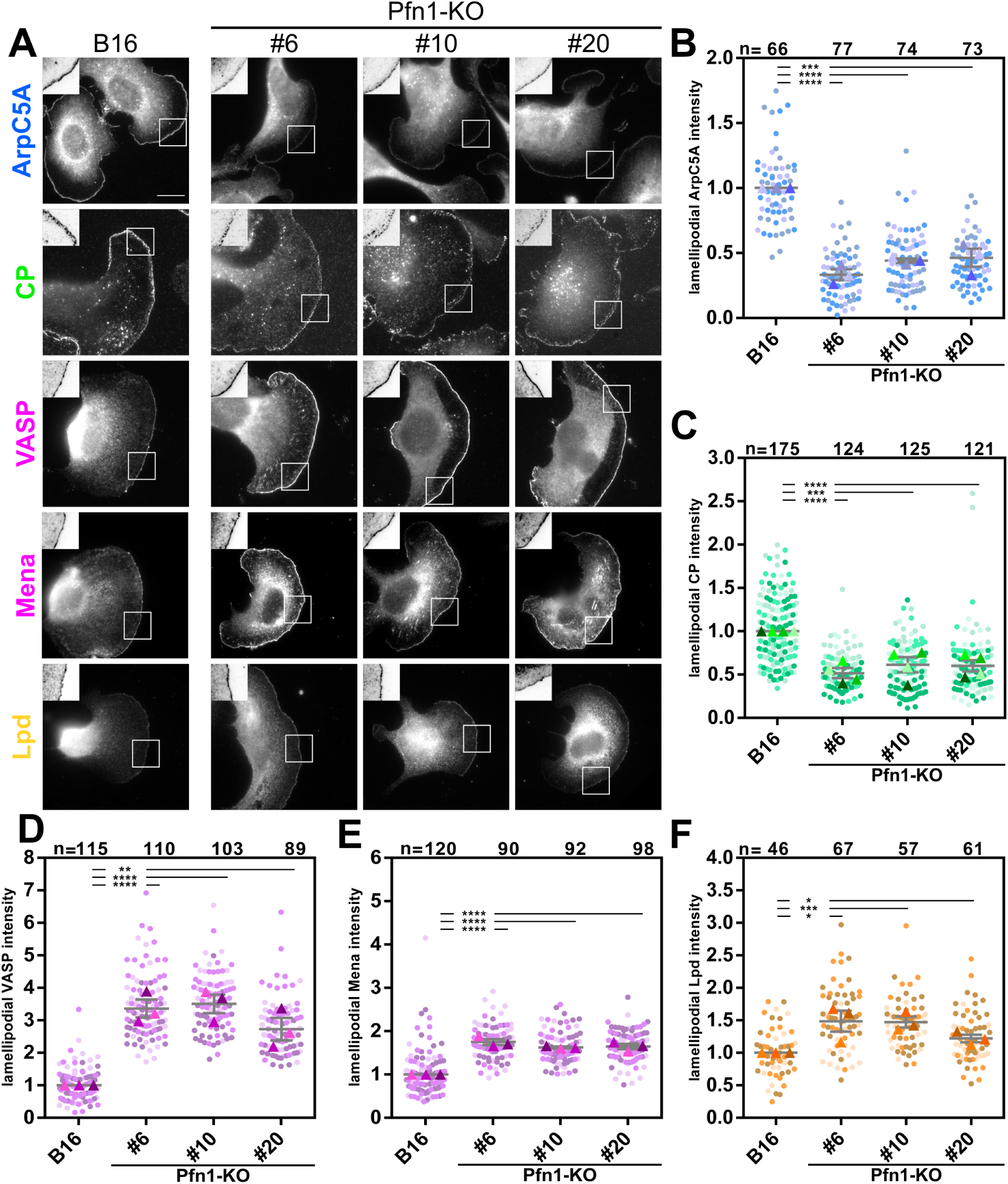
Consequences of profilin loss of function on lamellipodial accumulation of key actin assembly regulators. (**A**) Representative images of B16-F1 melanoma cells seeded on laminin and stained for Arp2/3 complex (ArpC5A, blue), capping protein (CP, green), Ena/VASP family members (VASP and Mena, pink), lamellipodin (Lpd, yellow). White squares depict insets displayed as magnified and inverted images at top left corners of each individual panel. Note respective changes in lamellipodial intensities that can be appreciated as black lines in inverted insets for each lamellipodial regulator. (**B**-**F**) Quantitations of lamellipodial fluorescence intensities of respective components, color-coded as described in A and displayed as superplots with independent experiments shown in distinct shadings of respective colors. Intensities measured of all components were normalized to the arithmetic mean of the WT population for each individual experiment (colored rectangles), n=measured cells from at least three independent experiments (four in C). For all data in B-F, datasets from each individual KO-line were statistically compared to WT using two sample two-sided T-tests;*p≤0.05; **p≤0.01; ***p≤0.005; ****p≤0.001.

Previous work on Pfn1-KO CAD cells suggested peripheral actin networks to be depleted of Arp2/3 complex incorporation entirely (Skruber *et al*., 2020). However, this finding appears inconsistent with both our previously published work on Arp2/3 complex subunit-KOs (Dimchev *et al*, 2021; Fassler *et al*, 2023) and the lamellipodia-like features of actin network protrusion and turnover observed in Pfn1-deficient cells (see above). We performed two experiments to confirm the contribution of Arp2/3 complex in the protrusion of Pfn1-deficient lamellipodia. First, we transiently expressed an EGFP-tagged version of the Arp2/3 complex subunit ArpC5L, revealing reduced (as compared to WT), but specific incorporation into the peripheral protrusions of Pfn1-KO clone #20 (Supplementary Video 6). Next, we used CRISPR/Cas9 to disrupt the essential Arp2 subunit of Arp2/3 complex, which is encoded by a single gene, and conducted detailed analyses on two distinct double-KO clones. If lamellipodia in Pfn1-KO cells were truly Arp2/3-independent, as suggested for Pfn1-deficient CAD cell protrusions (Skruber *et al*., 2020), the double-KO cells would be expected to display largely unaffected structures. However, we observed the opposite. Specifically, Pfn1-deficient B16-F1 clones additionally lacking Arp2 displayed an actin cytoskeleton mostly harboring prominent bundles at the periphery of concave edges, but lacking any obvious, protrusive actin network reminiscent of a lamellipodium (Extended Fig. 6A, B). Given that these cell clones are strongly impaired in both formin- and Arp2/3 complex-dependent actin assembly activities, it is not surprising that their migration was virtually eliminated compared to the parental Pfn1-KO or WT cells (Extended Fig. 6C, D, Supplementary Video 7).

We conclude that despite the marked reduction of Arp2/3 complex incorporation, the lamellipodia of Pfn1-deficient B16-F1 melanoma are strictly dependent on the Arp2/3 complex.

### Profilin suppression induces accumulation of Ena/VASP family members, which is crucial for effective lamellipodium protrusion and migration without Pfn-1

As opposed to Arp2/3 complex and CP, actin polymerases of the Ena/VASP family were strongly increased at the leading edges of lamellipodia in Pfn1-deficient cells (Fig. 2A), which could be quantitatively confirmed again in all three Pfn1-KO clones for the two independent family members VASP (Fig. 2D) and Mena (Fig. 2E). Although not essential for Ena/VASP family targeting to lamellipodia (Dimchev *et al*, 2020; Pokrant *et al*, 2023), the prominent Ena/VASP interactor lamellipodin (Lpd) (Colo *et al*, 2012; Law *et al*, 2013) was also significantly increased at these sites in Pfn1-deficient cells (Fig 2A, F).

Notably, VASP is well established to accumulate at lamellipodial edges in a fashion that positively correlates with their rate of protrusion, and hence of lamellipodial actin network polymerization (Krause *et al*, 2003; Rottner *et al*, 1999). In the absence of Pfn1, this rule appears to be broken, with VASP accumulation being enhanced in spite of reduced lamellipodial protrusion and actin polymerization, perhaps as compensation for suppression of Arp2/3-dependent actin assembly (see above) and coincident with downregulation of its competitor for barbed ends, CP (see also below). Interestingly, similar observations were made again in the Pfn1-KO CAD cell system, but the authors concluded that increased Ena/VASP family members at leading edges were non-functional in the absence of Pfn1 (Skruber *et al*., 2020). Furthermore, our previously-published data established that genetic disruption of all Ena/VASP family members increased the accumulation of both Arp2/3 complex and CP in lamellipodia (Damiano-Guercio *et al*, 2020), although the exact mechanism underlying this phenomenon remained unclear. In order to disentangle all these connections and activities, we analyzed cells in which Ena/VASP and Pfn1 were removed in combination. This approach allowed us to also test the previous hypothesis that accumulated Ena/VASP proteins are non-functional in the absence of profilin 1 (Skruber *et al*., 2020).

To this end, we started with Ena/VASP family triple-KO (EVM-KO) clone #23.7.66 (Damiano-Guercio *et al*., 2020), and disrupted the *Pfn1* gene as for wildtype cells above, again initially isolating three independently-generated, quadruple-KO cells (EVM+Pfn1-KO, clones #10, #19 and #27, Fig. 3A, B). Interestingly, cells lacking both Pfn1 and Ena/VASP family members were still able to form lamellipodia (Fig. 3B). However, these structures were significantly smaller, with further reduction of both lamellipodial width and actin filament mass compared to EVM-KO (Fig. 3C, D). We also analyzed lamellipodial protrusion rates in comparison to both WT as well as EVM- or Pfn1-KOs, and while this parameter was incrementally reduced from B16 WT to EVM- and Pfn1-KO, cells lacking both EVM and Pfn1 displayed an even more dramatic phenotype (Extended Fig. 7A, B). Lamellipodial actin assembly rates were also more dramatically affected in the absence of Pfn1 as compared to Ena/VASP, but slightly increased upon removal of both as compared to loss of Pfn1 alone, perhaps due to defective lamellipodial adhesion and hence increased rearward flow in these conditions (Extended Fig. 7C, D and unpublished data). Finally, the migration of EVM+Pfn1-KO clones was dramatically reduced compared to both EVM-KOs (Extended Fig. 7E, F, for migration tracks see extended Fig. 8) and to Pfn1-KO (Extended Fig. 7F, Supplementary Video 8). Collectively, the data demonstrate that Pfn1 loss-triggered Ena/VASP accumulation substantially contributes to the efficiency of lamellipodial protrusion and migration in the absence of Pfn1.

**Figure 3.**
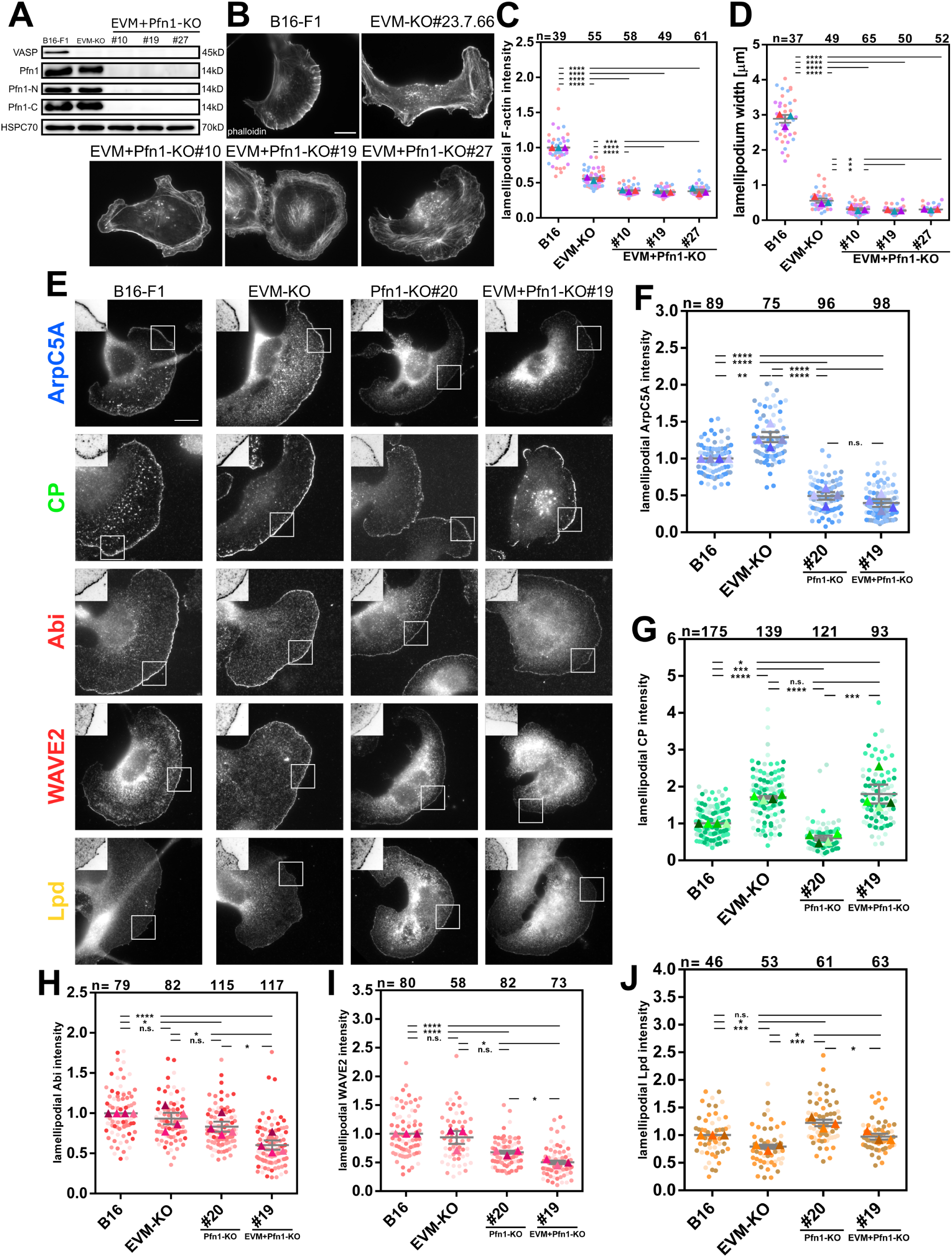
Profilin 1 removal in Ena/VASP-deficient cells compromises lamellipodia protrusion with simultaneous, differential impact on distinct core actin machinery components. (**A**) Western blotting of WT (B16-F1) as control and Ena/VASP-deficient cells before (EVM-KO) and after clonal isolation of cells treated with a Pfn1-specific guide construct. Following initial screening, quadruple-KO (EVM+Pfn1-KO) clones #10, #19 and #27 were selected for further analysis, and confirmed for the virtual absence of Pfn-1 using three independent antibodies specific for the N- and C-terminus or raised against the full-length protein, as indicated. Confirmatory VASP staining was employed for EVM-KO (Clone #23.7.66) and combined Pfn1-KOs, and HSP70 was used as loading control. (**B**) Representative cell images of the actin cytoskeleton stained with phalloidin of wildtype (B16-F1), EVM-KO and EVM+Pfn1-KO cell lines, as indicated. Note the incremental decrease of lamellipodial dimension (width) from WT to EVM-KO and EVM+Pfn1-KOs. The scale bar in the B16-F1 panel applies to all subpanels and equals 10µm. (**C**) Quantitation of lamellipodial F-actin intensities for the cell lines, shown in A, B. Data displayed as superplots as described for Fig. 1K. (**D**) Quantitation of lamellipodial widths in distinct cell lines, data as in Fig. 1L. EVM- and quadruple-KO datasets in C and D were statistically compared with B16 WT or EVM-KO, as indicated, using two sample two-sided T-tests; *p≤0.05; ***p≤0.005; ****p≤0.001. (**E**) Representative cell stainings of WT (B16-F1), EVM-KO and selected clones each for Pfn1-KO (clone #20) and EVM+Pfn1 (clone #19) for core actin machinery components as indicated. Color codes as described for Figure 2A, except that Ena/VASP family members were replaced by WRC subunits (Abi and WAVE2, red). (**F-J**) Quantitations of actin core machinery components, all assessed and displayed and color-coded as shown in E and described corresponding data in Fig. 2. n=measured cells from three (I, J) or four (F, G, H) independent experiments. For statistics, all datasets were compared with each other in all combinations using two sample two-sided T-tests; n.s., not significant, *p≤0.05; **p≤0.01; ***p≤0.005; ****p≤0.001.

### EVM removal upon Pfn1 loss of function has differential effects on Arp2/3 complex *versus* CP

As shown in Fig. 2, Arp2/3 complex and CP were strongly reduced in Pfn1-KO cell lines, but the opposite was previously observed in the B16-F1 clone lacking all EVM-family members (Damiano-Guercio *et al*., 2020), and employed above for generation of EVM+Pfn1 KO clones. Availability of the latter allowed asking whether Arp2/3 complex and CP accumulation would be experimentally inseparable or whether the mechanistic antagonism between Ena/VASP and CP would be dominant as compared to the co-association of Arp2/3 complex and CP. More specifically, the simultaneous removal of both Pfn1 and all Ena/VASP family members allowed distinguishing between the relative relevance of Pfn1 *versus* EVM for the extent of lamellipodial Arp2/3 complex incorporation. Indeed, the Pfn1 KO-induced suppression of Arp2/3 complex accumulation dominated over its Ena/VASP KO-triggered accumulation. In other words, the Ena/VASP removal-induced Arp2/3 complex enhancement observed previously (Damiano-Guercio *et al*., 2020) and confirmed here required the presence of Pfn-1, as Arp2/3 complex was equally suppressed following Pfn1 removal in lamellipodia harboring and lacking Ena/VASP proteins (Fig. 3E, F). The presence of Pfn1 thus dictates the extent of lamellipodial Arp2/3 complex incorporation irrespective of Ena/VASP, which is fascinating considering the key role for Pfn1 also in formin-mediated actin polymerization. Next, we performed the same type of analysis for lamellipodial CP, and to our surprise, the result was clear, but completely opposite to the behavior of Arp2/3 complex in quadruple-KO cells (Fig. 3E, G). This means that the EVM KO-induced increase of CP was preserved in the absence of Pfn1, or from another perspective, the Pfn1 KO-induced suppression of CP accumulation was reversed by additional Ena/VASP removal, positioning the increase of Ena/VASP accumulation upstream and hence causative of the reduction of CP accumulation in Pfn1-KO. To our knowledge, this is the first described experimental scenario, in which Arp2/3 complex and CP accumulation diverged, which shows that their incorporation into lamellipodial actin networks is functionally separable, in principle, and determined, at least in part, by distinct players among the actin-binding protein family. We have also assessed the accumulations in respective knockouts of the WRC subunits Abi and WAVE, but their accumulation patterns appeared poorly correlated with the changes observed in Arp2/3 complex accumulation. For instance, the increase in Arp2/3 complex levels observed in EVM-KO (Fig. 3F) was not accompanied by a corresponding increase in Abi or WAVE accumulation, both of which were virtually unchanged in intensity in these conditions (Fig. 3E, F), as seen previously (Damiano-Guercio *et al*., 2020). Furthermore, the marked suppression of Arp2/3 complex in Pfn1-KO was accompanied by at best a very modest reduction of WRC accumulation (Fig. 3H, I). And although in (EVM+Pfn1) quadruple KOs, Abi and WAVE were statistically significantly reduced as compared to Pfn1-KO alone, no significant change was observed between these two conditions for Arp2/3 complex incorporation (Fig. 3F). All this suggests that while WRC-mediated targeting and activation of Arp2/3 complex at the lamellipodium tip is essential for the formation of canonical lamellipodia (Dolati *et al*, 2018; Schaks *et al*., 2018), the quantitative changes observed here are hardly attributable to differences in WRC accumulation.

### Lpd is antagonistic to profilin concerning the lamellipodial accumulation of EVM *versus* CP

Quantification of Lpd at the tips of EVM-KO cell lamellipodia, which has not been explored previously to our knowledge, revealed a modest, but statistically significant decrease (Fig. 3E, J). This was quite surprising considering the common view of Lpd constituting an upstream regulator of Ena/VASP accumulation in lamellipodia (Law *et al*., 2013). Our data suggest that Ena/VASP and Lpd stabilize each other as modulators of protrusion in lamellipodia rather than Lpd serving as sole upstream recruiter of Ena/VASP (Law *et al*., 2013), also consistent with the fact that the latter could not be confirmed by various, comprehensive KO cell analyses (Dimchev *et al*., 2020; Pokrant *et al*., 2023 and this study). Furthermore, the increase of Lpd observed upon Pfn1 removal (see also above) was diminished in the absence of both Ena/VASP and Pfn1 (Fig. 3E, J), suggesting potential opposite effects for Ena/VASP *versus* Pfn1 on lamellipodial Lpd accumulation. To explore these potential, mutual antagonisms further, we performed the *vice versa* experiment, thus combined gene disruptions of Pfn1 and Lpd, and examined the effects on Ena/VASP and CP accumulation. Notably, it was impossible to genetically eliminate Pfn1 in the absence of Lpd entirely, but we suppressed Pfn1 expression in Lpd-KO#8 down to app. 20% of WT B16-F1 levels in six independent clones (Fig. 4A and data not shown). All six clones were explored in comparison to Lpd and Pfn1-KO alone for the accumulation of VASP (Fig. 4B, C). Interestingly, VASP that is modestly reduced in Lpd- and strongly increased in Pfn1-KO was found in intensities between WT and Pfn1-KO, as if the absence of Lpd partially, but statistically significantly counter-balanced the Pfn1 KO-induced increase in Ena-VASP. This was consistent for all six clones, largely independent of the amount of remaining Pfn1 expression (Fig. 4A). Due to the similarity of phenotypes, additional parameters were measured for two selected, representative Lpd+Pfn1-DKO clones only, #5 and #14. Interestingly, the significant changes in VASP accumulation (as compared to Pfn1-KO) did not in this case significantly impact on lamellipodial F-actin or Arp2/3 complex intensities (Fig. 4D, E, F), as Pfn1-suppression in the absence of Lpd did not decrease these intensities further as compared to cells lacking Pfn1 alone. In contrast, however, statistically significant differences were again observed for lamellipodial CP accumulation. In essence, all data for the latter were mirrored in a fashion opposite to Ena/VASP in distinct KO cell populations. CP accumulation increased upon Lpd removal (Fig. 4D, G), with concomitant decrease of VASP (Fig. 4B, C). Moreover, the Pfn1 KO-induced decrease of CP, which depended on the presence of Ena/VASP (see Fig 3E, G), was again partially reverted and hence alleviated in a statistically significant fashion in the absence of Lpd, so that the mean in each case was in between the levels measured for Pfn1-KO alone and WT. Based on aforementioned critical function of Ena/VASP on CP accumulation in the absence of profilin, we conclude that CP accumulation in respective combinations of knockouts followed in a mutually antagonistic fashion the accumulation of VASP, i.e. reduction of VASP increased CP and *vice versa*. At the same time, the respective biochemical activities on actin and Arp2/3 complex accumulation appeared to be balanced out as compared to Pfn1-KO alone (Fig. 4D, E, F).

**Figure 4.**
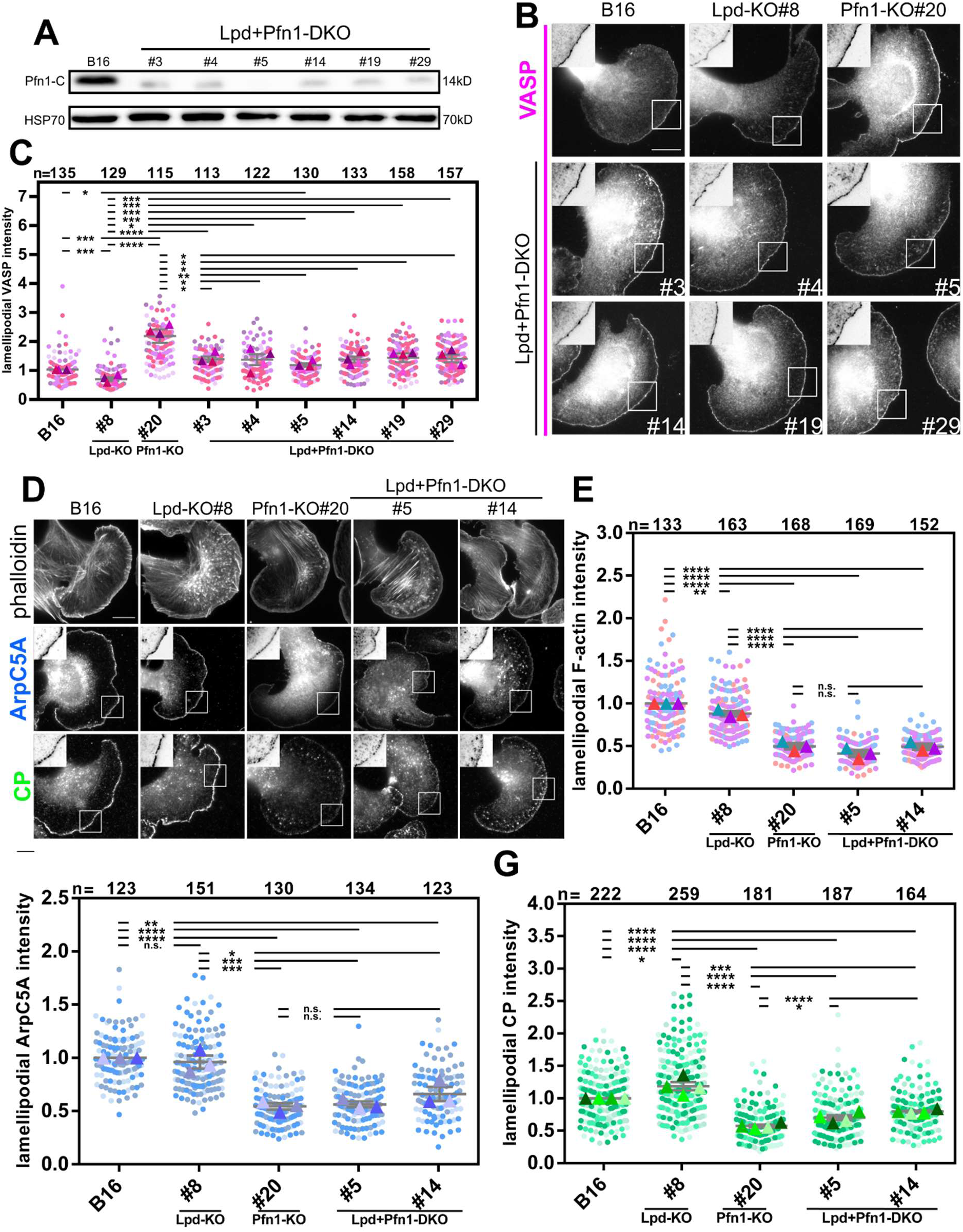
Suppression of profilin expression in the absence of lamellipodin generates intermediate phenotypes for VASP up-and CP downregulation in the lamellipodium. (**A**) Western blot displaying Pfn1-staining in cell extracts from lamellipodin (Lpd)-deficient lines isolated upon additional CRISPR/Cas9 targeting of the *Pfn1* gene, termed Lpd+Pfn1 double knockout (Lpd+Pfn1-DKO), as compared to B16-F1 control. Note that Pfn1 expression was reduced, but is still detectable in most (five out of six) DKO-clones. Furthermore, DKO-clone #5 lacking detectable staining here was confirmed to still harbor an allele with an in-frame deletion by TIDE-sequencing (not shown). DKO cell lines thus constituted Lpd-deficient cells with very low Pfn1 expression. (**B**) Immunolabeling using VASP-specific antibodies in B16-F1 wildtype (B16), Lpd-KO (clone #8), Pfn1-KO (clone #20) and Lpd1+Pfn1-DKO clones, as indicated (for clone numbers see right bottom corner). White squares depict insets displayed as magnified and inverted images at top left corners of each individual panel. Scale bar: 10 µm. (**C**) Quantitation of lamellipodial VASP intensities performed in a fashion identical to Fig. 2, but for cell lines as indicated. (**D**) Comparative stainings for the actin cytoskeleton (phalloidin), Arp2/3 complex (ArpC5A, blue) and capping protein (CP, green) in B16-F1 wildtype (B16), Lpd-KO#8, Pfn1-KO#20 and two representative Lpd+Pfn1-DKO (clones #5 and #14), as indicated. White squares in ArpC5A and CP panels depict regions inverted and magnified in top left corners as above. (**E** - **G**) Quantitation of lamellipodial F-actin, Arp2/3 complex (ArpC5A, blue) and capping protein (CP, green) in distinct cell lines as shown E. For statistics in C and E-G, all datasets were compared with each other in all combinations using two sample two-sided T-tests; n.s., not significant, *p≤0.05; **p≤0.01; ***p≤0.005; ****p≤0.001. In C, results from comparisons with WT B16 are shown only for one representative clone (double-KO#5), for the sake of clarity. All other DHO-clones differed from B16-F1 wildtype (B16) in a statistically significant fashion, with the exception of double-KO#4 (not shown).

To confirm the antagonistic relationship between Ena/VASP and CP in an independent, clean genetic system, we compared lamellipodial VASP and Mena accumulations in a cell line disrupted for CP (Funk *et al*., 2021). Indeed, complete removal of the CP β-subunit (CapZβ-KO) caused chaotic protrusion, but did not eliminate lamellipodia entirely (Funk *et al*., 2021; Hein *et al*., 2023), and this phenotype was accompanied by significant, lamellipodial tip accumulation of both VASP (Extended Fig. 6E, F) and Mena (Extended Fig. 6E, G). Interestingly, the extent of Ena/VASP accumulation in the complete absence of CP was even more dramatic than for cells examined in parallel and depleted of Pfn1 (clone #20) (Extended Fig. 6E, F, G), the latter of which are only reduced in, but not completely devoid of lamellipodial CP (see Fig. 2A, C). The resulting antagonism and additional mechanistic connections between all these key players of lamellipodial protrusion was further explored by mathematical modeling.

### Mathematical modelling reveals Ena/VASP and CP to compete for the same recruitment mechanism at the leading edge membrane, irrespective of barbed end binding

Mathematical modeling has been performed considering the interactions and biochemical activities of the key actin regulatory molecules in the lamellipodium studied above, including aside from actin and Arp2/3 complex WRC, CP and Ena/VASP. As described in detail in Extended Fig. 9, the model comprises the dynamic interactions of these molecules with the leading edge membrane, leading to a panel of variables, such as concentrations of WRC, CP or Ena/VASP at these sites, free or bound to filaments or (in case of WRC) to Arp2/3 or both (see Table 1 in Extended Fig. 9). The rates of these interactions, various concentrations of involved molecules as well as measured actin assembly rates in different KO cell lines are all parameters in our modeling (see Tables 2 and 4 in Extended Fig. 9). Pfn1 knockout enters the model as a reduction of branching rate. The values for these parameters have been determined by fitting the stationary states of the model to 14 criteria derived from the experimental results. These criteria constitute relative fluorescence intensity values of the model variables in different knockout conditions (see Table 3 in Extended Fig. 9). The parameter values meeting the criteria (allowing 20% deviation from means of experimental data) covered comparably broad ranges, suggesting the model to work robustly (Fig. M3 in Extended Fig. 9).

Key assumption in the model is that a reduction of polymerizable actin, as e.g. upon profilin suppression, is followed by reduced branching, as previously established (Carlsson, 2005; Carlsson *et al*, 2004). This view is fully recapitulated by our experimental data, as Pfn1 knockout not only reduces both lamellipodial actin (Fig. 1K) and Arp2/3 complex incorporation (Fig. 2A, B), but also actin expression (to app. 40%, Fig. 1G). Since profilin can catalyze the exchange of ATP for ADP on G-actin, amounts of polymerizable actin will likely be even lower, so were assumed in the model to constitute 20% of control (for details see Extended Fig. 9). The decrease of branching directly decreases F-actin and Arp2/3 complex densities in the lamellipodium. The density of WRC also decreases since less of it is involved in branching processes. Furthermore, the measured increase of EVM densities in Pfn1 knockout in spite of decreased lamellipodial F-actin and thus barbed ends demonstrates the EVM recruitment to the protrusion tip membrane to occur independently of barbed end density. The same argument applies to CP accumulation in the absence of Ena/VASP in spite of reduced actin filament and thus barbed end densities (Damiano-Guercio *et al*., 2020 and Fig. 3C, G). EVM and CP are thus recruited to the membrane independently of and likely prior to barbed end binding. Due to blocking polymerization, barbed end-bound CP is immediately advected away from the tip range by retrograde flow. In contrast, VASP can reside at the membrane upon barbed end binding as it drives filament elongation before eventually being removed if polymerization is stopped or too slow or, alternatively, its dissociation into the cytoplasm.

Our suggested mechanism explaining the experimental data implies that recruitment of EVM and CP to the same binding sites prior to capturing barbed ends continues upon Pfn-1 removal. However, with decreased barbed end densities in the virtual absence of profilin, the pool of recruiting binding sites fills up (see also Fig. 5). In accordance with the impact on the respective other component in EVM *versus* CP-KOs (Damiano-Guercio *et al*., 2020 and Extended Fig. 6E-G), we could fit the experimental data only by competitive binding of EVM and CP to these sites, and not just barbed ends. The fit of the model to experimental data generates rate constants for VASP *versus* CP binding such that EVM occupies a larger fraction of recruitment sites upon Pfn-1 removal. Finally, the model generated an even further increase of Ena/VASP accumulation if allowing solutions with CP leading edge levels in Pfn1-KO lower than the aforementioned 20% deviation (Figs. M5-7 in Extended Fig. 9). All this confirms the strict mutual antagonism between Ena/VASP and CP accumulation at the lamellipodial tip membrane, and can be viewed as *in silico* counterpart of the experimental data shown in Fig. 4. Again, this mechanism only works if VASP and CP compete for the same recruitment sites at the membrane instead of just barbed ends as common for historical models of protrusion regulation (Bear & Gertler, 2009). How this idea of competitive EVM *versus* CP recruitment will be reconciled with the proposed functions of some of their direct or indirect interactors, such as Lpd or WRC in case of EVM (Colo *et al*., 2012; Kage *et al*, 2022) *versus* CARMIL for CP (Fujiwara *et al*, 2014; Mooren *et al*, 2024) will be exciting topics of future investigations.

**Figure 5.**
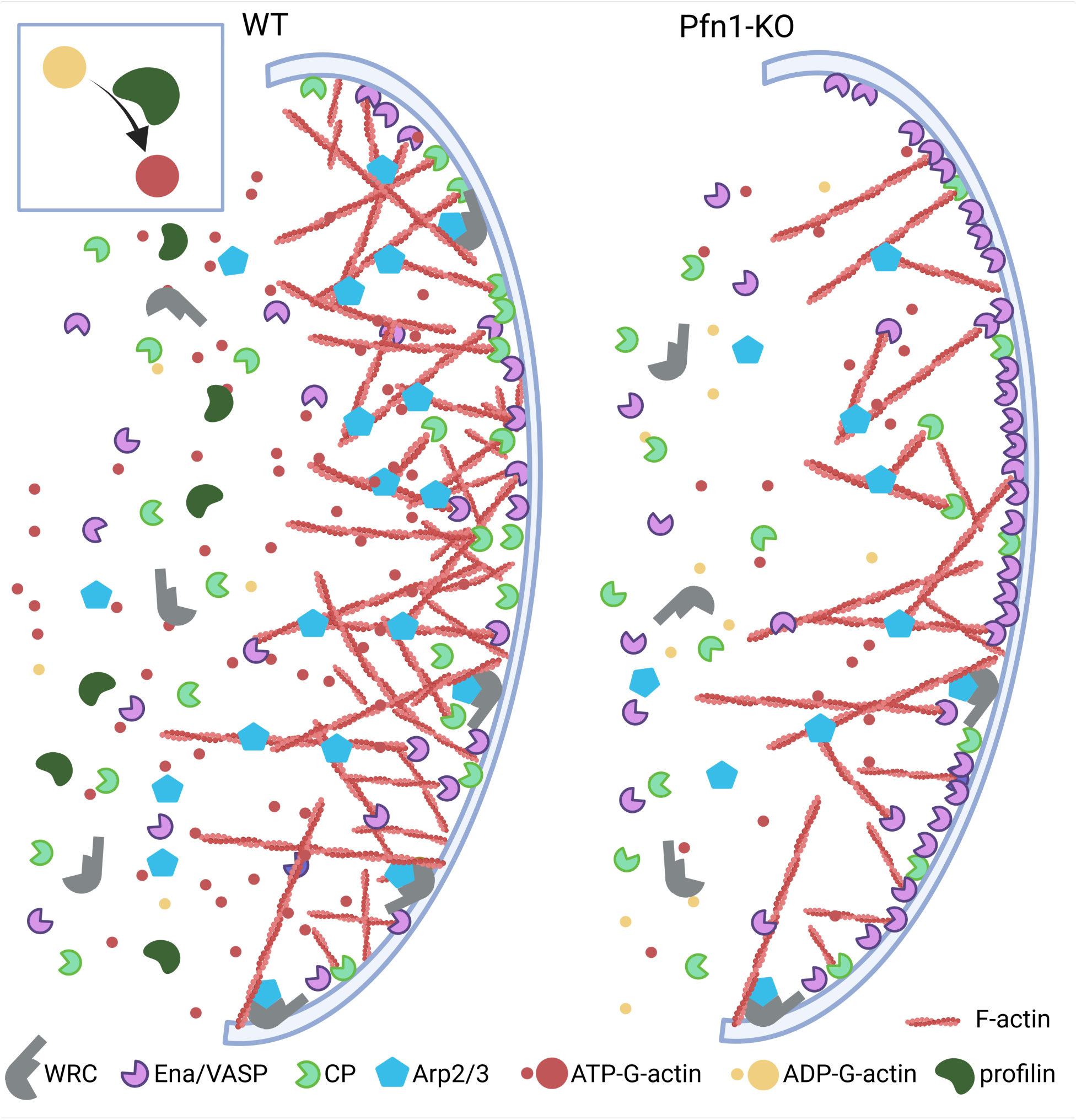
Summary of morphological and molecular transformations caused by profilin 1 (Pfn-1) removal. Profilin loss of function reduces total actin levels reflected by both reduced polymerizable actin monomer and lamellipodial actin filaments and coinciding with strongly suppressed, but not eliminated, WRC-dependent Arp2/3 complex incorporation. In contrast, these transformations effect enhanced accumulation at lamellipodial edges of Ena/VASP family members and, as a consequence, reduced CP accumulation. This hypothesis for effects on the actin core machinery upon profilin loss of function is recapitulated by mathematical modeling (see Extended Fig. 9). Inset on top left emphasizes nucleotide exchange on actin by profilin, symbols for actin regulators are depicted at the bottom and not drawn to scale.

## Discussion

Dynamic actin filament remodeling, such as in the lamellipodium, involves several widely conserved factors, whose function remain incompletely understood despite extensive study. This is particularly true for core regulators like profilin (Pfn), Ena/VASP family polymerases, and heterodimeric capping protein (CP). Here, we investigate their complex interrelationships within Arp2/3 complex-dependent lamellipodial protrusions. Historically, Arp2/3 complex-mediated actin nucleation and branching were considered antagonistic to formin-dependent elongation, which relies on Pfn. Thus, Arp2/3-generated branched networks and formin-driven linear filaments were viewed as antagonizing structures competing for a limited pool of polymerizable actin (Suarez & Kovar, 2016). However, our findings, along with previous studies, challenge this oversimplified view, besides the fact that Arp2/3 complex is now also recognized to generate linear filaments downstream of SPIN90, not just branches (Cao *et al*, 2023; Skruber *et al*., 2020).

Using cell lines with minimal Pfn expression, we confirm its essential role in formin-dependent actin structures, such as filopodia triggered by mDia2 or FMNL3. Surprisingly, Pfn also enhances Arp2/3 complex activity in lamellipodia, promoting actin branching. Pfn depletion reduces, but does not eliminate, Arp2/3 complex incorporation into lamellipodia, likely due to a severe drop in polymerizable actin, partly driven by a ∼60% decrease in total actin levels. The transformation of lamellipodia in the virtual absence of Pfn includes a prominent accumulation of Ena/VASP family polymerases and a concomitant suppression of its antagonist, CP. Disrupting both Ena/VASP and Pfn1 reveals that Ena/VASP controls CP but not Arp2/3 incorporation, since the latter remains low even when Ena/VASP is removed in Pfn1-KO cells. Thus, while VASP antagonizes CP, Pfn promotes Arp2/3 complex activity, contrary to previous assumptions (Rotty *et al*., 2015; Suarez *et al*., 2015).

These findings highlight that each of these factors do not solely influence each other indirectly through tuning actin assembly dynamics, but through direct reciprocal effects. For instance, Pfn1 depletion-induced VASP accumulation is partly reversed by suppressing its interactor Lpd. This in turn alleviates the CP reduction seen with Pfn1 loss alone, supporting a direct mutually antagonistic relationship between Ena/VASP and CP in lamellipodia.

Our mathematical modeling robustly recapitulates these observations in Pfn1- or combined Pfn1- and Ena/VASP-KO cell lines. The model shows that the antagonism in VASP *versus* CP accumulation is best explained by additional, antagonistic membrane targeting rather than sole direct competition for barbed ends. Furthermore, obtained modeling solutions represented broad parameter ranges, illustrating the robustness of our framework in describing actin network remodeling.

The model also provides novel quantitative insights into the relationships between involved factors and/or their interactions (see scaled variables in Figs. M4 and M7 in Extended Fig. 9). One key prediction is the relatively rare interaction between WRC and Arp2/3 complex (∼1% of free WRC molecules at the membrane), similar to the low frequency of WRC/Arp2/3 capturing a filament. Despite being crucial for lamellipodia assembly (Dimchev *et al*., 2021; Suraneni *et al*, 2012; Wu *et al*, 2012; Mietkowska et al., unpublished data), Arp2/3-mediated branching is far less frequent than elongation events. Moreover, filament-bound WRC levels are comparable to or slightly higher than free WRCs, suggesting a significant role for the complex in barbed end elongation as distributive polymerase (Bieling *et al*, 2018). Yet, VASP-bound filaments outnumber WRC-bound ones by up to tenfold, emphasizing the dominance of molecular players solely engaged in elongation. Interestingly, free VASP at the membrane is even more abundant than filament-bound VASP, matching CP levels and explaining why unbound filaments are rare (Figs. M4 and M7 in Extended Fig. 9).

All these data suggest that lamellipodial actin filament generation and elongation are tightly coordinated across its core regulators. However, the molecular basis of the Ena/VASP and Arp2/3 complex antagonism remains unclear. Since WRC biases Arp2/3 complex activation towards the membrane, Ena/VASP-bound filaments may be suboptimal substrates for branching. Additionally, the precise mechanism underlying Ena/VASP and CP competition for leading edge targeting remains unknown, though our experiments and modeling suggest it must be largely independent of barbed end binding. Notably, our previous BioID data indicate that Ena/VASP and CP exist in close proximity (<10nm) (Pokrant *et al*., 2023), suggesting potential indirect interactions within larger protein complexes. Future work will aim to resolve these outstanding questions.

## Materials and methods

### Cell culture and transfections

B16-F1 mouse melanoma cells (purchased from American Type Culture Collection, Manassas, VA, CRL-6323) were cultivated according to standard tissue culture conditions. In brief, cells were grown at 37°C in an atmosphere containing 7.5% CO2 in Dulbecco’s modified Eagle medium (DMEM) containing 4.5 g/l glucose (Life Technologies, Thermo Fisher Scientific, Germany) and 10% fetal calf serum (FCS; Gibco, Paisley, UK), 2 mM glutamine and 1% penicillin-streptomycin (Thermo Fisher Scientific). Wildtype B16-F1 cells were splitted every other day at densities at least 1:10 and 1:20, and distinct knockout cell lines described below adapted for splitting dilutions according to their respective growth capabilities. Previously published CRISPR-KO cell lines employed for analyses and/or used as parental lines for additional gene disruptions were EVM-KO (clone #23.7.66) (Damiano-Guercio *et al*., 2020) and Lpd-KO (clone #8) (Dimchev *et al*., 2020).

For transfections of cells with respective DNA vectors, JetPrime transfection reagent (Polyplus Transfection, Illkirch, France) was used according to the manufacturer’s instructions. For microscopy experiments, B16-F1 cells were seeded onto glass coverslips pre-coated for 1 h at room temperature with 25 ng/μl laminin (L-2020; Sigma-Aldrich, Germany) in coating buffer (50 mM Tris-HCl, pH 7.4, and 150 mM NaCl) if not stated otherwise. Constructs for live cell fluorescence imaging of actin cytoskeleton dynamics were EGFP-tagged, human β-actin (PT3265-5, Clontech Inc., Mountain View, CA, USA), mDia2-FL and -ΔDAD (Block *et al*, 2008) as well as ArpC5B (Rottner *et al*, 2006).

*Dictyostelium discoideum* AX2-214 wildtype (WT) and PfnI/II-deficient cells (Haugwitz *et al*., 1994) were cultivated at 21° C in HL5-C medium containing glucose (HLC0102, Formedium) supplemented with 50 µg/ml penicillin and 40 µg/ml streptomycin (Carl Roth) on polystyrene-coated petri dishes. For homologous recombination, cells were transfected by electroporation as described below.

### Gene disruptions in B16-F1 melanoma and *D. discoideum*

Gene disruptions by CRISPR/Cas9-mediated genome editing in B16-F1 melanoma followed by single cell cloning and expansion were performed essentially as described before (Kage *et al*, 2017). Specifically, disruption of the gene locus encoding profilin 1 (*Pfn1*) in each cell line (B16-F1 wildtype, EVM-KO#23.7.66, Lpd-KO#8 or Pfn2-KO#41, for generation of the latter see below) was done by targeting exon 1 and using guide-sequence 5’- TGTCAGGACGCGGCCATCGT-3’, cloned into CRISPR expression vector pSpCas9(BB)-2A-Puro (Addgene 48138). Successful knockout clones were initially screened by Western blotting followed by Sanger sequencing of genomic DNA amplicons subcloned into bacterial colonies (for Pfn1 KO-lines #6, #10 and #20) or by TIDE-sequencing in case of EVM+Pfn1-KOs and Lpd+Pfn1-KOs (data not shown). Specifically, for genomic DNA extractions prior to sequencing, cells were pelleted and incubated at 55℃ overnight in lysis buffer (100 mM Tris-HCl pH 8.5, 5 mM EDTA, 0.2% SDS, 200 mM NaCl) containing 40 μg/ml proteinase K, followed by nucleic acid extraction by phenol/chloroform precipitation. The targeted locus was PCR-amplified with forward and reverse primers 5’-ATCTGAAGCTTAGCTAACAGAGCCGCGTTC-3’ and 5’- ATCTGCTCGAGAGGGTCTAAGGCCGGTCAC-3’, respectively. In case of Pfn1 single KOs, amplicons were purified with a NucleoSpin Gel and a PCR clean-up kit following manufacturer’s instructions (Macherey&Nagel), inserted into a Zero Blunt TOPO vector using a commercial Zero Blunt TOPO Cloning Kit (Invitrogen) and transformed into competent bacteria. Upon inoculation and growth of single bacterial colonies, derived plasmid DNAs were purified using a NucleoSpin Plasmid kit (Macherey&Nagel) and sequenced by Eurofin Genomics (Ebersberg, Germany) using sequencing primer 5’-CAGGAAACAGCTATGAC-3’. Knockout lines obtained upon targeting of the *Pfn1*-locus in EVM-KO or Lpd-KO cells were sequenced by TIDE-sequencing (Brinkman *et al*, 2014). Clones with mutations exclusively causing frameshifts on all detected alleles were selected for further characterization.

The attempt to disrupt Pfn1 in the Pfn2-KO background (clone #41) in Extended Fig. 2C-E was not followed by sequencing, as no reduction of Pfn1 expression upon subcloning could be detected by Western Blotting (Extended Fig. 2E). The stable Pfn2-KO (clone #41) used for this experiment was originally generated as follows: Exon 1 of the *Pfn2* gene was targeted using guide-sequence 5’-GTCTGGGCAGCCACGGCCGG-3’, again cloned into pSpCas9(BB)-2A-Puro vector (Addgene 48139) and transfected into B16-F1 wildtype cells for stable KO generation essentially as described for the *Pfn1* locus above, except that Lipofectamine® 2000 Transfection Reagent (Invitrogen) was used following manufacturer’s instructions. Upon clone selection by Western blotting, PCR fragments from the amplified target regions of each clone were sub-cloned into pcDNA3 vector, followed by Sanger sequencing of individual colonies. Clone #41 turned out to lack any wildtype allele and was thus selected for further experiments.

The gene locus giving rise to expression of the Arp2 subunit of Arp2/3 complex (*Actr2*) was targeted in Pfn1-KO (clone #20) using 5’-TCCTTCCAGTTTGTGAAGTG-3’ as guide-sequence. Upon editing and selection of promising clones by Western blotting, the selected region within exon 2 was PCR-amplified with oligonucleotides 5’-TAAAAGCCAGGGTAAGCAAGGC-3’ and 5’-TTCTGACCCTCCCTTTGCTTTT-3’ as forward and reverse primers, respectively, and analyzed by TIDE-sequencing.

Disruption of the *proC* gene in *D. discoideum* as shown in Extended Fig. 2F, G was performed as follows: The targeting vector was constructed by amplification of a PstI/BamHI 425-base pair 5′-fragment and a SalI/HindIII 553-base pair 3′-fragment flanking the *proC* gene. Amplified fragments were subsequently inserted into the corresponding restriction enzyme sites of plasmid pLPBLP containing a blasticidin S resistance cassette (Faix *et al*, 2004). The resulting vector was cleaved with BamHI and SalI, and used to disrupt the *proC* gene encoding PfnIII. Primers sequences for the 5′ fragment were 5′- CGCGGATCCATCTGATTGATTTATATTG-3′ and 5′- GATCTGCAGCTTGCCAAGTCATTATTTC -3′, and for the 3′ fragment 5′- GAGAAGCTTCCATATGTATAATATCTCCTTC-3′ and 5′-TATGTCGACGTTTTCGAAATTTTCTATAATCTTG-3′. The fidelity of the targeting vector was verified by sequencing. The PfnIII-targeting construct was then transfected into WT or PfnI/II-KO cells by electroporation using the Xcell Gene Pulser (Bio-Rad) following preset protocol 4-2-6. Transfected cells were selected 24 h after transfection with 10 μg/ml blasticidin (Invivogen) for seven days and then maintained in growth medium in the absence of blasticidin. Null-mutants were identified by diagnostic PCRs of genomic DNA as described previously (Faix *et al*., 2004). The primers used for analysis were AU: 5′-GTTTCGATCTGGTTGTGAATGGAATGG-3′, AD: 5’-ACATATAAAAATTCTCGCGTTAAATACG-3’, AD2: 5’-CAAAGGTGATACCCAATCAACTG-3’ and Bsr: 5’-CAAACAGTTACTCGTCCTATATACG-3’.

### Immunoblotting and quantitation of Pfn2 or actin expression levels

For preparation of whole cell extracts for immunoblotting, cells were washed once with PBS followed by treatment with lysis buffer (2% SDS, 10% glycerol, 63mM Tris-HCl pH 6.8) and sonicated to shear genomic DNA. Upon addition of β-mercaptoethanol and bromphenolblue to obtain Laemmli buffer, all lysates were heated for 15 min at 95°C and stored at −20°C before usage. To determine expression levels of respective proteins or confirm CRISPR/Cas9- mediated gene disruptions, proteins were separated by SDS-PAGE and transferred onto PVDF membranes for immunoblotting following standard procedures. Primary antibodies were as follows: homemade, polyclonal anti-profilin1 raised against the full length protein (Pfn1, 1:1000, kindly provided by Prof. Dr. Walter Witke, University of Bonn) (Witke *et al*, 1998); polyclonal anti-Pfn1 raised against the N-terminus (Pfn1-N, P7749, 1:3000, Sigma-Aldrich); polyclonal anti-Pfn1 raised against the C-terminus (Pfn1-C, P7624, 1:3000, Sigma-Aldrich); monoclonal anti-Pfn2 (sc-100955, 1:200, Santa Cruz Animal Health); homemade, polyclonal anti-VASP (5500-A, 1:2000) (Damiano-Guercio *et al*., 2020); rabbit monoclonal anti-Arp2 (ab129018, 1:2000, Abcam); monoclonal anti-GAPDH (MAB374, 1:10000, MERCK); monoclonal anti-HSP70 (also known as HSPA8/HSC70, sc-7298, 1:10000, Santa Cruz Animal Health). HRP-conjugated secondary antibodies used were goat anti-mouse IgG (H+L) (111-035-062, 1:10000, Dianova) or goat-anti-rabbit IgG (H+L) (111-035-045, 1:10000, Dianova). Chemiluminescence obtained upon incubation with ECL Prime Western Blotting Detection Reagent (GE Healthcare) was recorded using an ECL Chemocam imager (Intas, Germany).

For determination of Pfn2 expression levels, quantitative Western blots were performed essentially as described (Funk *et al*., 2019). Briefly, serial dilutions of recombinant profilin2 were run alongside TCA-precipitated lysates from B16-F1 wild-type and profilin1 knockout cells on 15% SDS-PAGE gels. Proteins were transferred to PVDF membranes (Merck) using Tris-Glycine buffer. Membranes were blocked with 5% milk in TBS-T (TBS + 0.2% Tween20) for 1 hr at RT, then incubated with anti-profilin2 antibody (1:500, #sc-100955, Santa Cruz). Infrared-labeled donkey anti-mouse secondary antibodies (1:10,000, #925–32212, #926–68073, LI-COR) were used for detection on a LI-COR Odyssey CLx. Fluorescence intensities were analyzed using ImageJ. Equal-sized rectangular areas were selected for each protein band. Intensity profiles were generated using the “Plot Lanes” function, and background signal was subtracted by adjacent background regions. Signal intensities, represented as the area under the intensity profiles, were integrated using the tracing tool. A standard curve was generated by plotting integrated intensities against reference profilin2 protein amounts and fitting a linear function. This standard curve was used to calculate profilin2 mass per cell. The concentration of profilin2 was then calculated as follows:

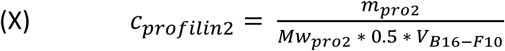

with m_pro2_ being the mass of profilin2 per cell, Mw_pro2_ being the molecular weight of mouse profilin2 (15032.21 g/mol) and V_B16-F10_ being the total cell volume of B16-F10 cells, which we assumed to be similar to B16-F1. We factored in only half of the total cellular volume because profilin is largely excluded from organelles that occupy roughly 50% of the cell volume.

For quantification of global actin levels, proteins in total cell lysates were separated by SDS-PAGE and blotted onto nitrocellulose. The blots were then either probed with a rabbit polyclonal pan anti-actin antibody (Hein *et al*., 2023) (1:1000) or an anti-β-tubulin antibody (MAB1864, Merck Millipore, 1:500) overnight. Primary antibodies in immunoblots were visualized by enhanced chemiluminescence (ECL) using peroxidase-coupled anti-rabbit IgG (111-035-003, Dianova, 1:10,000) or anti-mouse IgG (115-035-062, Dianova, 1:10,000) using the ChemiDoc MP Imaging System driven by Image Lab software (Biorad, Hercules, CA, USA). After background subtraction of 16-bit images, band intensities of actin were normalized to respective tubulin signals, and relative protein levels calculated in Pfn1-KO mutants compared with B16-F1 control (WT).

For determination of F-/G-actin ratios in B16-F1 WT and Pfn1-deficient cells, nearly confluent cultures, corresponding to about 5 x 10^6^ cells, were first rinsed twice with ice-cold PBS. Cells from each plate were detached from the plate with a cell scraper using 300 µL of cold lysis buffer containing 20mM Hepes, pH 7.2, 100mM NaCl, 10mM KCl, 2mM MgCl_2_, 5mM KPO_4_, 5mM EGTA, 5mM ATP, 2mM DTT, 5% sucrose, 0.5% Triton X-100, 0.5% NP-40, 5mM benzamidine (434760, Sigma-Aldrich), 0.1mM AEBSF hydrochloride (A1421, Applichem) and 0.75µM phalloidin (2141, Sigma-Aldrich). The crude lysates were then transferred into 1.5 mL reaction tubes and placed onto a rotary shaker for further homogenization at 4°C. After 30 min, 200µL samples were centrifuged at 150,000 x g for 1h using an Optima Max-XP tabletop ultracentrifuge (Beckman Instruments, Palo Alto, CA, USA). Subsequently, the samples of the supernatant and the pellet fractions were brought to 300µL with SDS sample buffer, and aliquots subjected to SDS-PAGE followed by immunoblotting. The blots were then probed with aforementioned rabbit polyclonal pan anti-actin antibody and visualized by ECL using peroxidase-coupled anti-rabbit IgG as described above. After background subtraction of 16-bit images, actin band intensities were determined densitometrically using ImageJ software, and ratios of actin levels in pellet and supernatant fractions calculated in Pfn1-KO mutants *versus* B16-F1 controls.

### Phalloidin stainings, immunolabelings and quantitations of lamellipodial features

For immunolabelings or phalloidin stainings, cells were seeded onto glass coverslips coated with laminin as described above and allowed to spread overnight. Both were done using standard procedures essentially as described in detail before (Kage *et al*., 2022).

For phalloidin stainings, cells were fixed for 20 min with a mixture of pre-warmed (37°C) paraformaldehyde (PFA, 4% in PBS) and 0.25% glutaraldehyde, followed by permeabilization in 0.05% Triton-X 100/PBS for 40 sec. Fixed and permeabilized samples were then extensively washed and subjected to phalloidin-staining (Atto 488-labelled, AD488-81 or Atto 550-labelled, AD 550-8, stock diluted 1:50 into PBS, ATTO-TEC GmbH, Siegen, Germany). DNA was visualized with 40,6-diamidino-2-phenylindole (DAPI) (Sigma-Aldrich, 1:1000). This was followed by embedding and image acquisition. For immunolabelings, cells were treated in an identical fashion, except that fixation was done with 4% PFA alone and that the staining procedure was initiated with 5% horse serum dissolved in 1% BSA/PBS followed by primary and secondary antibodies diluted in 1% BSA/PBS, except for primary undiluted supernatants. Additional exceptions were anti-VASP and anti-CP stainings, for which a mixture of 4% PFA and picric acid (added to app. 0.7‰) and 3% glyoxal at pH 5.0 (Richter *et al*, 2018) were used, respectively. Primary antibodies were: Anti-Arp2/3 complex (anti-ArpC5A, clone 323H3, homemade, undiluted supernatant) (Olazabal *et al*, 2002); polyclonal anti-Mena, -VASP and - WAVE2 used at 1:100, 1:400 and 1:1000 dilutions, respectively (Damiano-Guercio *et al*., 2020); anti-Lpd (3917, 1:200) (Krause *et al*, 2004); monoclonal anti-capping protein α1/α2 (mAb B5 12.3, 1:4, Developmental Hybridoma Bank, University of Iowa, IA, USA); monoclonal anti-Abi (W8.3, hybridoma supernatant, 1:20) (Innocenti *et al*, 2004). Secondary antibodies were Alexa Fluor^TM^ 594-conjugated Goat anti-Mouse IgG (H+L) (A11032, ThermoFisher Scientific) and Goat anti-Rabbit IgG (H+L) (A11037, ThermoFisher Scientific) used at concentrations according to manufacturer’s instructions. Image acquisition using epifluorescence microscopy was routinely done on an inverted microscope (Axiovert 100TV or Axio Observer, Carl Zeiss, Jena) equipped with 1.3NA 40x or 1.4NA 63x and 100x objectives, widefield illumination (HXP120, Zeiss or DG4, Sutter Instrument) and a Coolsnap-HQ2 camera (Photometrics) controlled by Metamorph (Molecular Devices) or VisiView® software (Visitron Systems GmbH, Puchheim, Germany). Images shown in Ext. data Fig. 3 were captured with a Zeiss confocal LSM 980 with Airyscan 2 equipped with a Plan-Apochromat 63×/1.46 NA oil DIC or C-Apochromat 40×/1.20 W Korr objective using 405 and 561-nm laser lines. Emitted light was detected in the wavelength ranges of 410 to 473 and 576 to 610, respectively.

Quantitations of lamellipodial F-actin or other lamellipodial factors were done as follows: For F-actin, average pixel intensities of three randomly-chosen lamellipodial regions from phalloidin-stained samples lacking microspikes were measured followed by subtraction of a nearby, extracellular region defined as background (for representative regions, see Figure 1J). The arithmetic mean of these three regions per cell corresponded to single data-point in measurements displayed as superplot data, as e.g. in Fig. 1K. All other lamellipodial components were measured in an analogous fashion, but considering their specific localizations, i.e. localization throughout the network (in case of F-actin or Arp2/3 complex) *versus* lamellipodial tip components (e.g. VASP, WRC, Lpd). Microspike intensities were quantified in an analogous fashion, but now measuring respective fluorescence intensities throughout microspike bundles.

Lamellipodial width was quantified from phalloidin-stained images with MetaMorph by drawing a line from the lamellipodium tip to the distal edge of the lamella, followed by measuring line lengths. Again, three randomly chosen, lamellipodial sections were measured, averaged and expressed as single data point/cell in μm. Microspike lengths were determined and displayed in analogous fashion.

### Random cell migration, high magnification video microscopy and fluorescence recovery after photobleaching (FRAP)

Random cell migration assays were performed precisely as described before (Kage *et al*., 2022), and data analyzed in ImageJ (https://imagej.net/ij/) using the Manual Tracking Plugin as well as the Chemotaxis and Migration Tool (Ibidi).

High magnification time-lapse phase-contrast and fluorescence video microscopy was done on an Axio Observer (Carl Zeiss, Jena) equipped with a DG4 (Sutter Instrument) for fluorescence and a VIS-LED for phase-contrast illumination, a 1.4NA 100x Ph3 oil immersion objective and a back-illuminated, cooled, charge-coupled-device camera (CoolSnap HQ2, Photometrics), all driven by VisiView® Software (Visitron Systems GmbH, Puchheim, Germany). Kymography followed by protrusion rate determination were done as described before (Schaks *et al*., 2018), and data displayed as superplots.

For FRAP experiments, bleaching was done on the same system with a 405nm diode laser at 60mW output power to ensure rapid, near-complete bleaching of EGFP-actin in the lamellipodium. Fluorescence recovery of the lamellipodial network was then recorded at 0.5 Hertz and an exposure time of 300ms. Rates of lamellipodial network assembly *versus* protrusion and rearward flow were manually read out aided again by kymography.

### Image processing, statistics and illustrations

Adjustments for brightness and contrast were performed using Metamorph software (Molecular Devices) or ImageJ. Image and data analysis was mostly done with Metamorph, ImageJ or Fiji, and Figures were assembled with Graphpad Prism 8 and Inkscape. All quantitative datasets were routinely repeated at least three independent times, and shown as arithmetic means ± SEMs (error bars), unless indicated otherwise. Statistics were done using SigmaPlot 12.0 (Systat Software, Erkrath, Germany) or Origin 2021 (OriginLab), and presented as provided in respective Figure legends. The schemes in Figs 5 and M1 (Extended Fig. 9) were drawn using BioRender (www.biorender.com).

## Mathematical modeling

The model formulates our hypotheses as a system of differential equations subsequently fitted to criteria defined by experiments, as described in Extended Fig. 9.

## Acknowledgments

The authors thank Brigitte Denker, Katja Mummenbrauer and Svenja Koch for expert technical assistance. We would also like to thank Profs Walter Witke (University of Bonn, Germany) and Giorgio Scita (IFOM Milan, Italy) as well as Dr. Matthias Krause (King’s College London, UK) for kindly sharing antibodies.

This work was supported in part by the German Research Council (DFG), individual grants RO 2414/8-1 (to K.R.), Fa330/12-3 (to J.F.), BI 1998/2-1 (to P.B.) and Fa350/19-1 (to M.F.), intramural core grants from the Helmholtz Society (to T.E.B.S. and K.R.) and by the Carl Trygger Foundation (R.K.). C.L. and Y.T. are thankful for stipends granted by the Life-Science Foundation (LSS, Munich) and China Scholarship Council (CSC), respectively.

## Author contributions

Y.T., M.S., M.M., J.S., R.B.N., S.K. and Z.L. performed experiments, interpreted data and/or made and edited Figures. C.L., T.E.B.S. and R.K. interpreted data and/or provided essential reagents. P.B. and J.F. performed experiments, interpreted data and/or made Figures. K.R. and M.F. developed the project together, with K.R. being responsible for wet experiments and M.F. for mathematical modeling. K.R. and M.F. wrote large parts of the paper with conceptual help from P.B., T.E.B.S. and J.F. All authors contributed to text corrections and proofreading.

## SUPPLEMENTARY INFORMATION TO

### Supplementary Figure legends

**Extended Figure 1.**
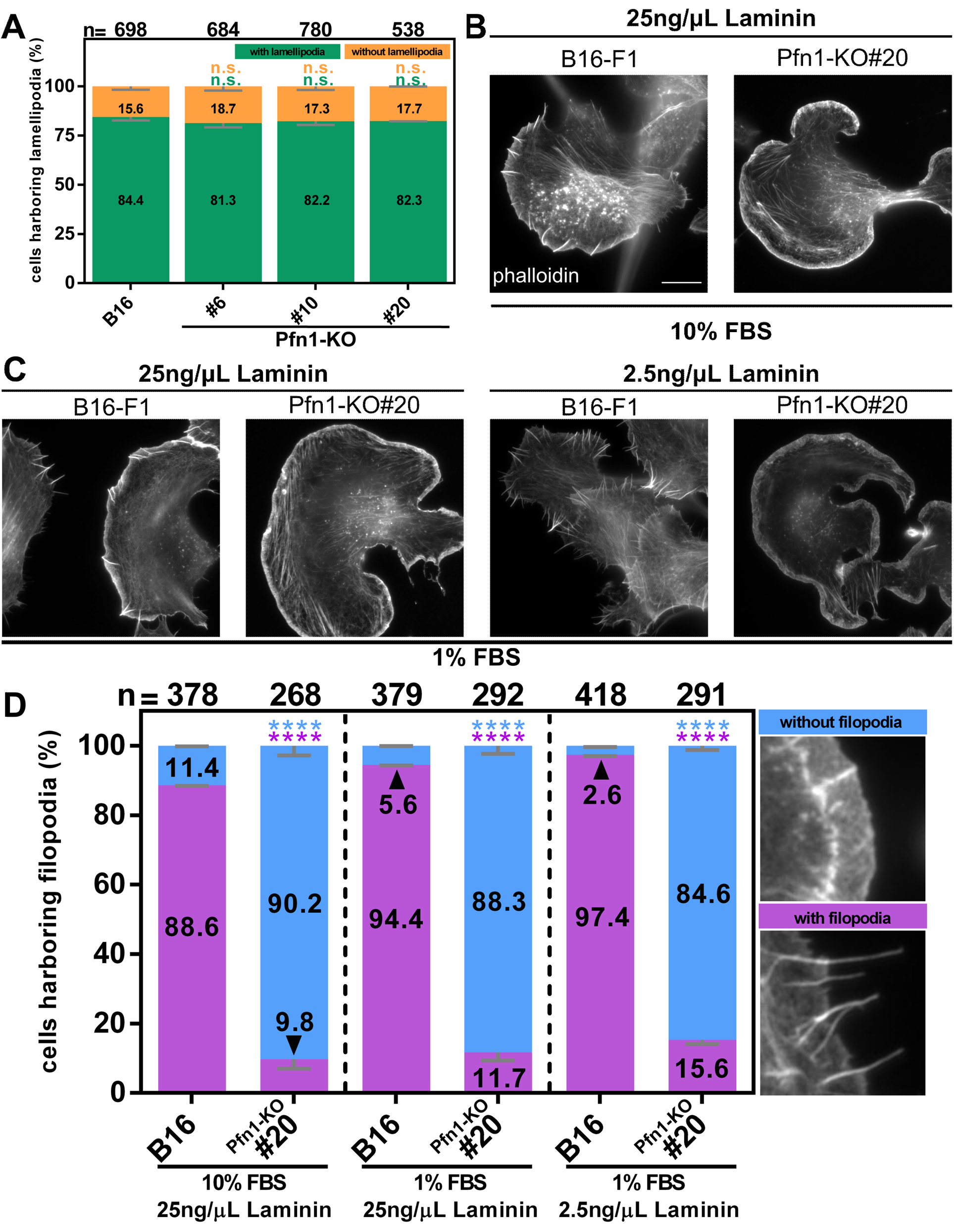
Profilin loss of function abrogates the frequency of filopodia, but not lamellipodia formation. (**A**) Quantitation of the percentage of cells capable of forming lamellipodia, irrespective of their size, precise morphological appearance or dimension - see color-code for cells with *versus* without lamellipodia at top right. Bars and numbers show arithmetic means and standard errors of means (SEMs) from three independent experiments, n=numbers of individual cells analyzed. No statistically significant differences in the frequency of lamellipodia formation in individual Pfn1-KO cell samples as compared to B16-F1 wildtype (B16) were observed. Two-sided two sample T-test, n.s. not significant. (**B**) Representative images of wildtype (B16-F1) and Pfn1-KO (clone #20) cells fixed and stained for the actin cytoskeleton using fluorescent phalloidin upon seeding on laminin (25ng/µl) in full growth medium (10% FBS), which constitutes our standard conditions for migration and lamellipodia formation. In these conditions, most (but not all) peripheral actin bundles appear as microspikes remaining embedded into lamellipodia in control cells, which are largely missing, however, in Pfn1-KO cells (see independent quantitation of microspike numbers in Fig. 1M). (**C**) Images of WT (B16-F1) and Pfn1-KO (clone #20) cells growing in conditions suboptimal for lamellipodia formation (reduced FBS and/or laminin concentrations, as indicated) and thus known to enhance extent and frequency of microspike or filopodia formation. Left panels display cells in 1% FBS and common laminin (25ng/µl) concentration and right panels 1% FBS and 2.5ng/ml laminin concentration. Note the explosive formation of filopodia and microspikes in the latter condition in wildtype cells (B16-F1), but very little response in this respect in the representative Pfn1-KO cell. (**D**) Quantitation of filopodia formation in all aforementioned conditions; for color-code of cells with versus without filopodia see panels on the right. In spite of the large qualitative and quantitative differences in filopodia formation for individual cells in the different conditions in B and C, the sole frequency of filopodia formation was incrementally, but only modestly increased in all conditions. However, albeit incremental increases in filopodia formation towards enhanced filopodia formation in low serum and laminin concentrations in both genotypes, filopodia formation was drastically suppressed in all conditions in the absence of Pfn-1. Statistically significant reduction of filopodia formation in the absence of Pfn1 in each case was confirmed by two-sided two sample T-test, ****p≤0.001.

**Extended Figure 2.**
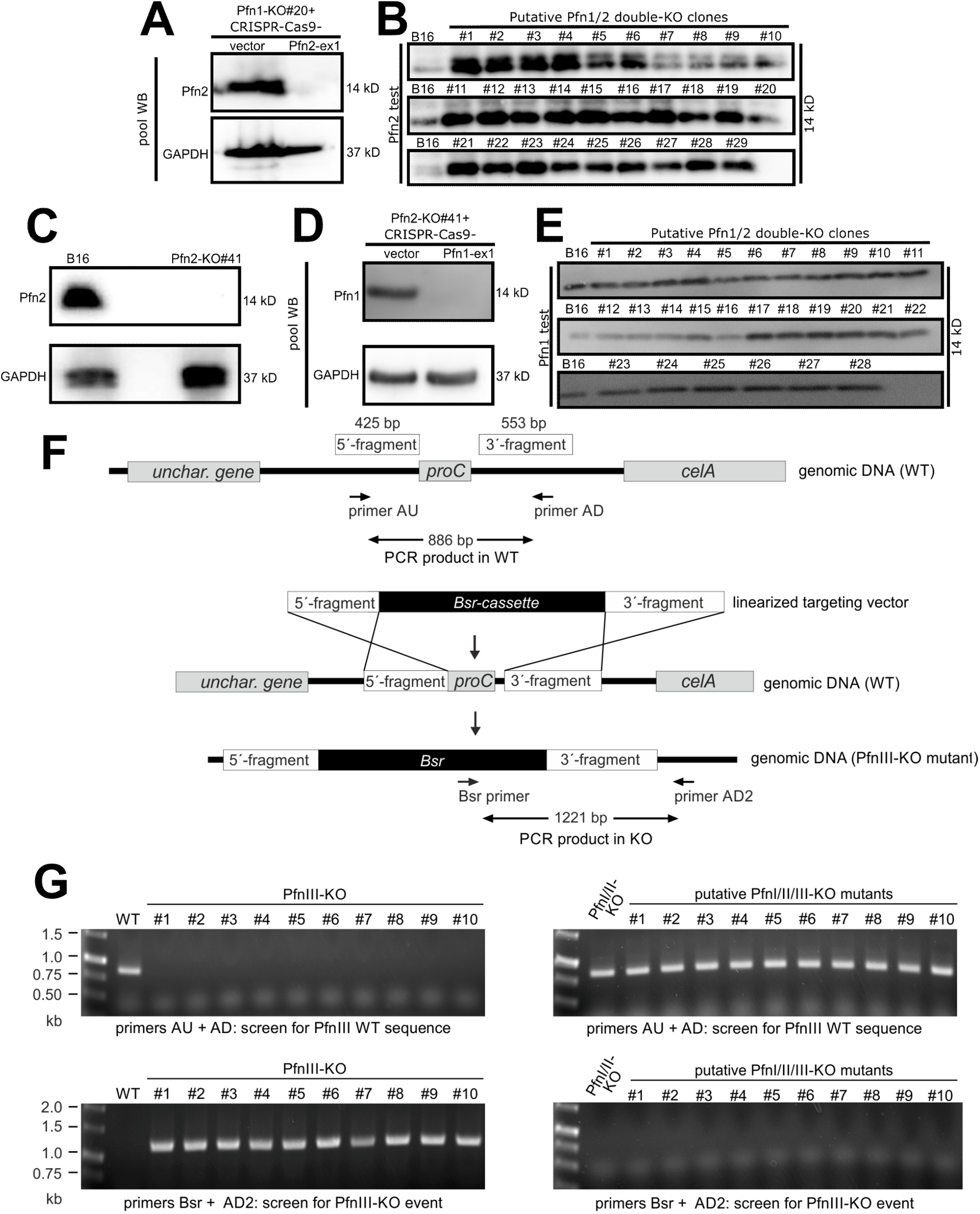
Consecutive genetic disruption of Pfn1 and −2 in mammalian or PfnI, -II and -III in *D. discoideum* cells does not lead to viable cell progeny. (**A**) Expression of Pfn2 in Pfn1-KO (clone #20) cells before and after acute disruption of the gene by CRISPR, using a Pfn2-specific guide sequence targeting exon 1 (Pfn2-ex1). Note the efficient suppression of Pfn2 expression app. 7 days after transient transfection with Pfn2-specific CRISPR-construct and selection for transfected cells with puromycin. (**B**) Screening Western blot with the aim to select for Pfn2-deficient cell line upon single cell cloning of the cell population shown in A (right). Note that out of 29 single-cell clones of Pfn1-KO#20 isolated and expanded upon acute Pfn2 disruption shown in A, not a single one of them lacked Pfn2expression. (**C**) Western blot showing the absence of Pfn2 in a B16-F1 clone selected upon CRISPR/Cas9 targeting of Pfn2. (**D**) Expression of Pfn1 in Pfn2-KO (clone #41) cells before and after acute disruption of the gene by CRISPR, using the Pfn1-specific guide targeting exon 1 (Pfn1-ex1). Again, note the efficiency of suppression of Pfn1 expression app. 7 days after transient transfection with Pfn1-specific CRISPR-construct and selection for transfected cells with puromycin. (**E**) Screening Western blot to select for stably Pfn1-depleted lines in the Pfn2-KO background upon single cell cloning of the cell population shown in D (right). 28 clones were analyzed, but all showed Pfn1 expression, indicating that, similar to the results shown in subpanels A and B, clones devoid of both Pfn1 and -2 are not viable or capable of proper cell growth and thus lost during the subcloning procedure. (**F**, **G**) PfnIII can be efficiently disrupted in *Dictyostelium* Ax2 WT cells, but not in PFNI/II-KO cells. (**F**) Schematic representation of the organization of the *proC* gene encoding profilinIII (PfnIII), construction of the targeting vector and generation of PfnIII- O cells. The 5′ and 3′ non-coding sequences flanking the *proC* gene, as indicated, were inserted into pLPBLP containing the Bsr resistance cassette. The linear targeting vector was then used to disrupt the *proC* gene by homologous recombination in Ax2WT and PfnI/II-KO mutant cells. (**G**) Upon homologous recombination, Bsr-resistant cells were examined for the presence of an intact or disrupted PfnIII allele by PCR employing two different primer sets. Top panels: the primer combination AU and AD as shown in F specifically identifies the presence of an intact wild-type allele. Bottom panels: the primer combination Bsr and AD2 as shown in F identifies insertion of the targeting vector into the *proC* gene and successful disruption of the gene. The high efficiency with which PfnIII can be eliminated in WT cells and the inability to inactivate PfnIII in the PfnI/II-KO mutant background strongly suggest an essential function of PfnIII in *D. discoideum* cells.

**Extended Figure 3.**
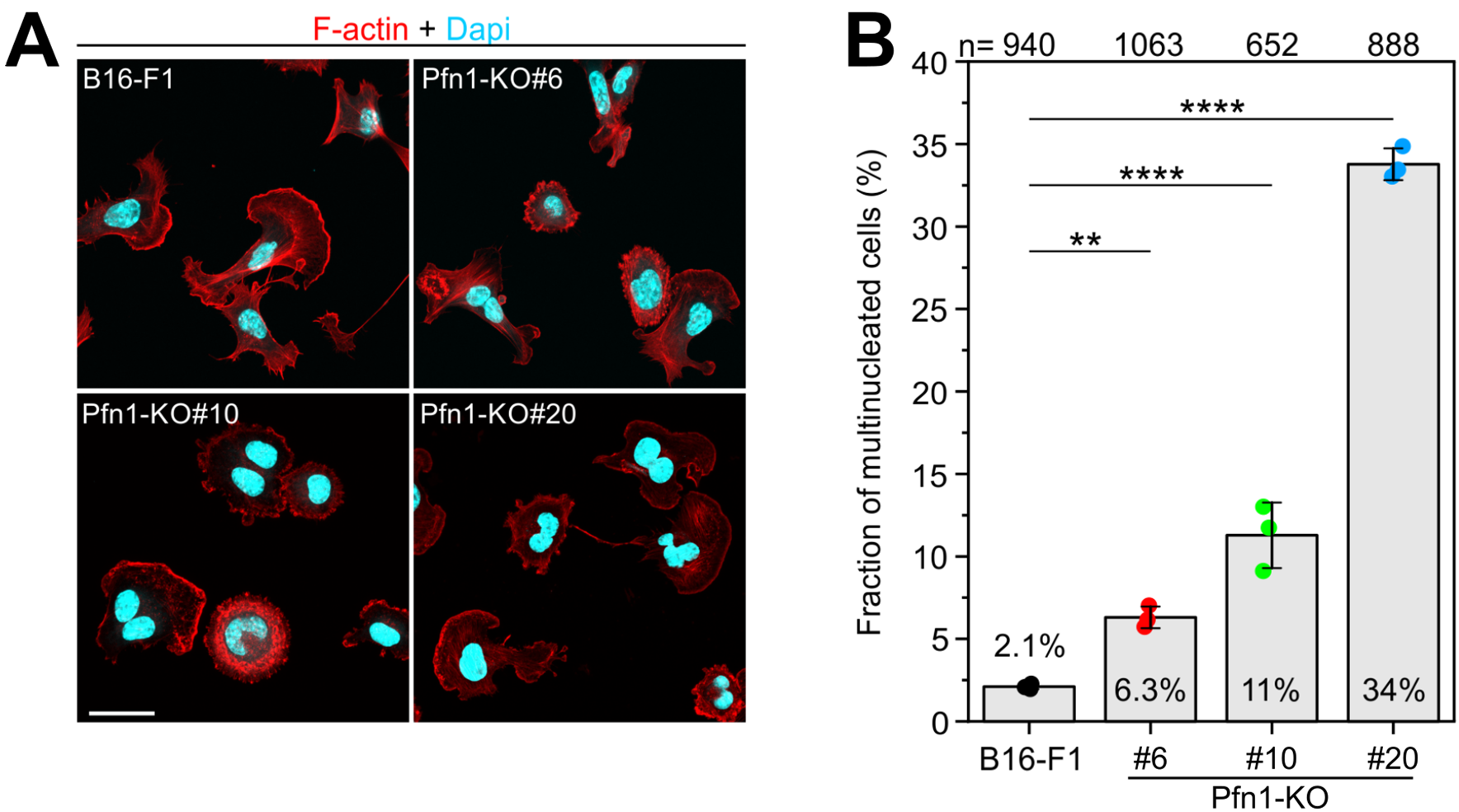
Pfn1-KO cell lines display varying degrees of multinucleation that correlate with remaining levels of Pfn2. (**A**) Representative images of WT (B16-F1) and Pfn1-KO cell lines (#6, #10 and #20), as indicated, fixed and double-stained with phalloidin (red) for the actin cytoskeleton and DAPI (turquois) for nuclear DNA. Scale bar, 20µm. (**B**) Quantification of the fraction of multinucleated WT and mutant cells as shown in A. Note the incremental increase of multinucleation in Pfn1-KOs, negatively correlating with the residual amounts of Pfn2, and hence overall profilin activity present in these lines (compare Fig. 1E). Bars and error bars represent arithmetic means and standard deviations from three independent experiments (N, colored dots). n, number of cells analyzed. One-way ANOVA and Tukey Multiple Comparison test were used to reveal statistically significant differences between datasets. **p≤0.01; ****p≤0.001.

**Extended Figure 4.**
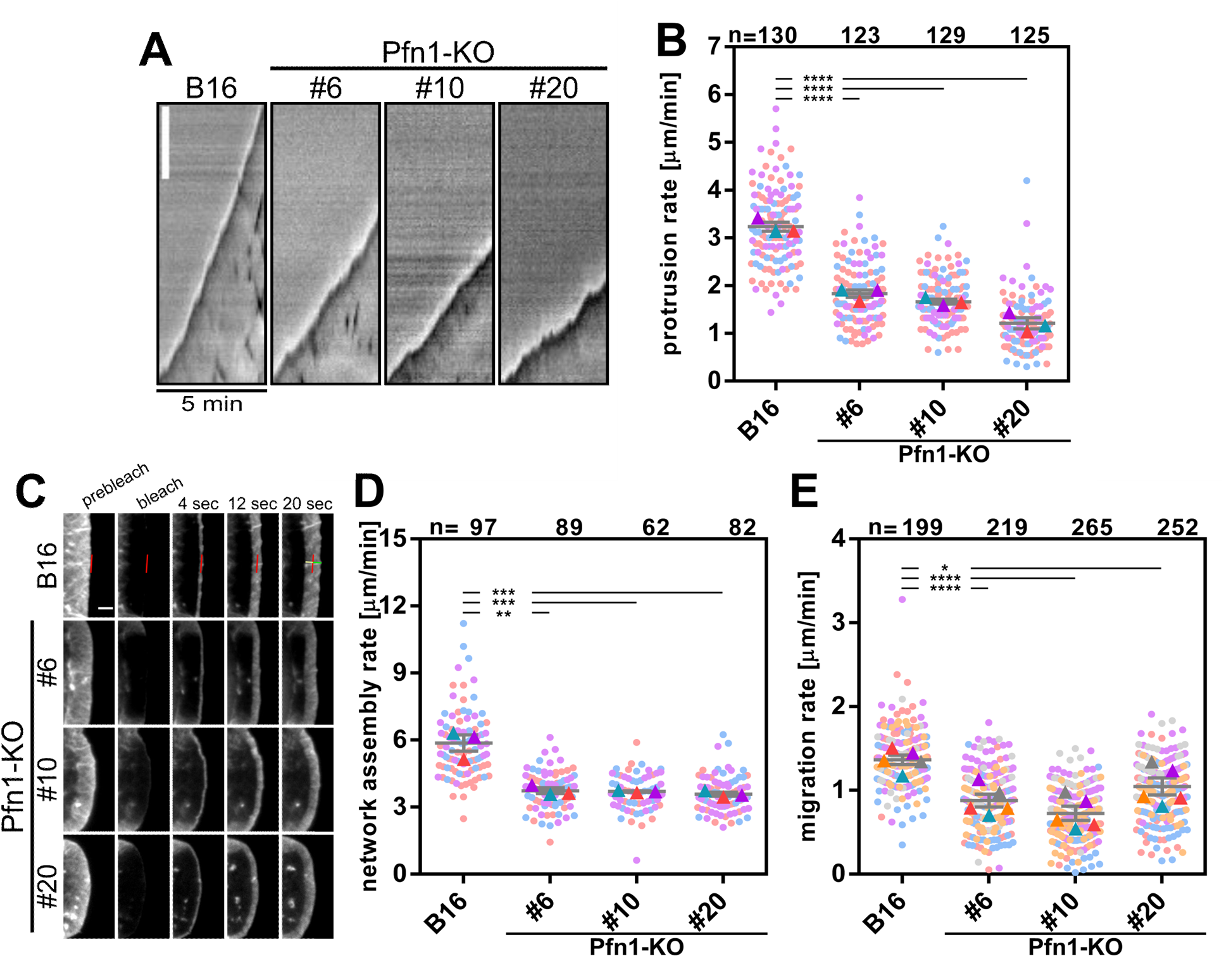
Pfn1-deficient cells display reduced lamellipodial protrusion, actin network assembly and overall random cell migration. (**A**) Kymographs (space-time plots) of representative lamellipodial edges of cells with genotypes as indicated and recorded by time-lapse phase contrast microscopy; time is as provided and white scale bar corresponds to 3µm. (**B**) Quantitation of lamellipodial protrusion rates determined from kymographs of individual lamellipodia (shown in A) for the different cell types. The protrusion rates determined for each cell line in three independent experiments are shown as so called superplots, displaying measurements from individual cells and their arithmetic means as color-coded dots and triangles, respectively. Grey lines and error bars display means and standard errors of the means (SEMs), n = numbers of measured cells. Datasets from each individual KO-line were statistically compared to WT (B16) using two sample two-sided T-tests; ****p≤0.001. (**C**) Fluorescence recovery after photobleaching (FRAP) analysis in lamellipodia of cells transiently expressing EGFP-tagged actin. Fluorescence recovery from the front following the bleach is taken as direct readout of lamellipodium network assembly rate. This equals the dimension of the lamellipodial actin network building up over time (20 seconds in this case, as indicated, see B16 panels) and constitutes the sum of protrusion distance (green line in front of the red line marking the lamellipodial edge in the prebleach frame) and rearward flow distance (yellow distance proximal to the prebleach red line). Scale bar, 2µm. For the movie corresponding to these data, see Supplementary Video 5. (**D**) Quantitation of network assembly rate assessed as representatively shown in C. Data are displayed as superplots in analogy to what’s described for protrusion rates in B. Statistics: two sample two-sided T-tests; **p≤0.01; ***p≤0.005. (**E**) Quantitation of random migration rates of B16-F1 wildtype (B16) and Pfn1-KO cell lines as indicated. Migration rates were extracted from migration tracks (see below) over time periods of 10 hours. Data are displayed as superplot data as above, but from five independent experiments (distinct colors). For a compiled representative movie including all cell lines see Supplementary Video 2. Statistics: two sample two-sided T-tests; *p≤0.05; ****p≤0.001.

**Extended Figure 5.**
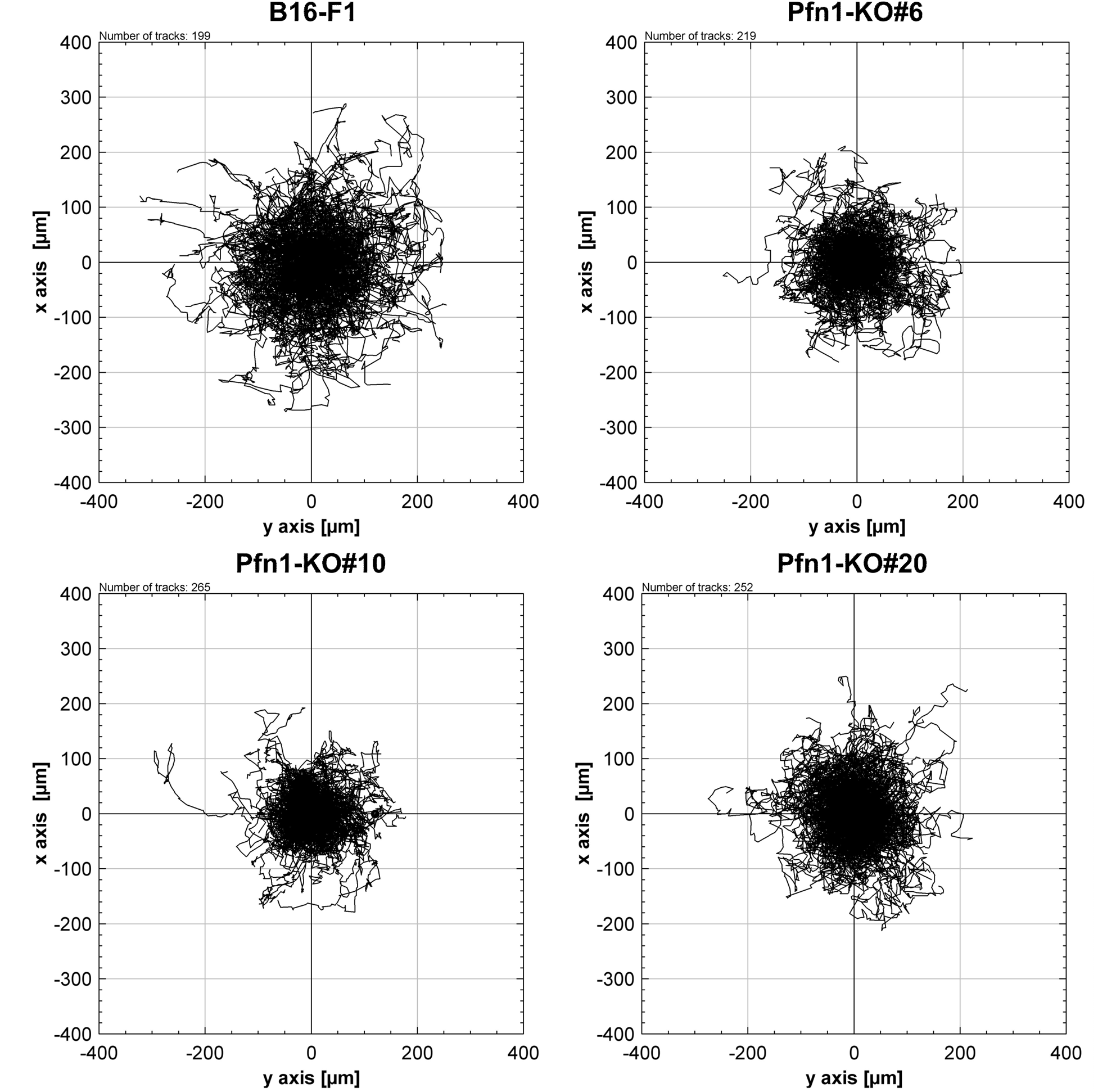
Pfn1-deficient B16-F1 melanoma cells migrate shorter distances than WT cells. Trajectory plots as obtained from random migration movies of distinct cell lines migrating on laminin and exemplarily shown in Supplementary Video 2. Each line corresponds to an individually tracked cell, the starting point of which was positioned in the center of the coordinate system. Cells were tracked for a maximum of 10 hours. Note the modest size reduction of trajectory plots as compared to wildtype in lines lacking Pfn1, consistent with the reduced migration rates (Extended Fig. 4E) as well as reduced lamellipodial protrusion rate (Extended Fig. 4B). This fits the notion that efficient migration of B16-F1 melanoma on laminin strongly correlates with the capability of lamellipodia formation (see Kage *et al*, 2022; Schaks *et al*, 2018).

**Extended Figure 6.**
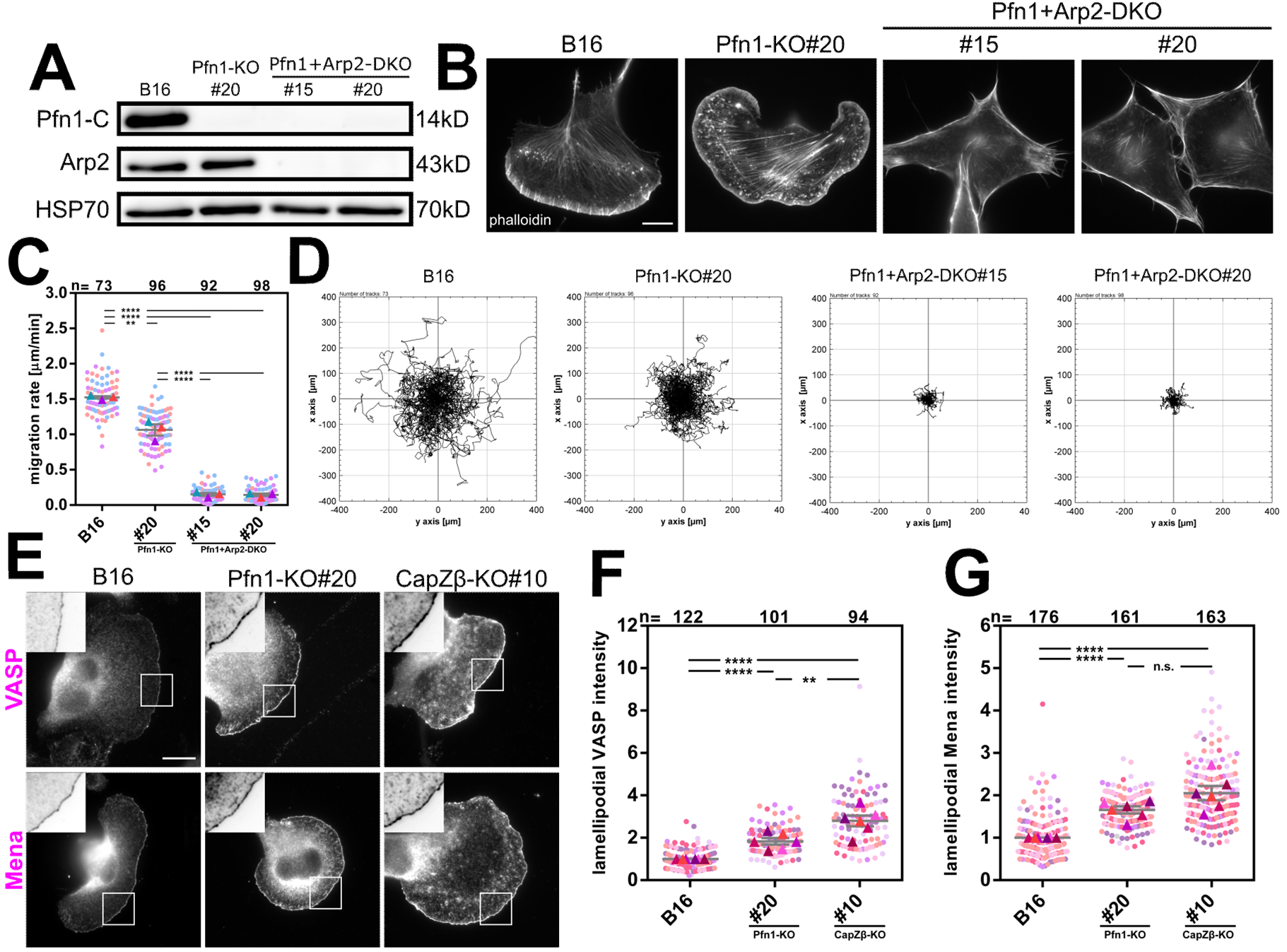
Pfn1-deficient lamellipodia require Arp2/3 complex, and the Pfn1-KO induced upregulation of lamellipodial Ena/VASP family members is phenocopied in CP-KO. (**A**) Western blot showing the removal of Arp2 in Pfn1-KO (clone #20). Two independent Pfn1+Arp2 double-KO (DKO) clones were selected, as indicated, and subjected to further analyses. Note the complete absence of detectable Arp2 staining, which coincided with the absence of any functional allele encoding Arp2 (data not shown). (**B**) B16-F1 cells with genotype, as indicated, upon growth and fixation on laminin-coated coverslips, and stained for the actin cytoskeleton with fluorescently-labeled phalloidin. Note the complete abrogation of Pfn1-deficient lamellipodia in the absence of the obligatory Arp2/3 complex subunit Arp2. In these cells, intracellular actin filament accumulation is largely compromised and cell edges featured by prominent, peripheral actin bundles, without any discernible signs of lamellipodia formation, as expected (Dimchev *et al*, 2021; Suraneni *et al*, 2012; Wu *et al*, 2012). Scale bar, 10µm. (**C**) Quantitation of random cell migration in B16-F1 wildtype (B16) as compared to the parental Pfn1-KO#20 and the same cells upon disruption of the Arp2 gene. Note the virtual elimination of migration in cells lacking both Arp2/3 complex activity and Pfn1, the latter in essence translating into lost processive formin activity (see above). (**D**) Trajectory plots obtained from the random migration movies of distinct cell lines migrating on laminin and used for the quantitation in C, for representative movies, see also Supplementary Video 7. As before, each track corresponds to an individual cell, the starting point of which was positioned in the center of the coordinate system. Cells were tracked for a maximum of 10 hours. (**E** - **G**) Comparison and quantitation of Ena/VASP protein accumulation at the lamellipodium tip in B16-F1 wildtype (WT) *versus* Pfn1-KO (clone #20) and a B16-F1 cell line disrupted for heterodimeric CP (CapZβ-KO#10) (see Funk *et al*, 2021). Representative stainings of cells of respective genotype are shown in E, as indicated. White rectangles in subpanels in E correspond to insets magnified and inverted at the respective top left of each panel, illustrating the increase in accumulation at the lamellipodium edge in Pfn1- and even more so CP-KO cells. Scale bar, 10µm. For quantitations in F and G, intensities measured for VASP and Mena were normalized to the arithmetic mean of the WT population in each individual experiment (colored rectangles), n=measured cells from at least six independent experiments. All datasets in F and G were statistically compared with each other using two sample two-sided T-tests; n.s. not significant; **p≤0.01; ****p≤0.001.

**Extended Figure 7.**
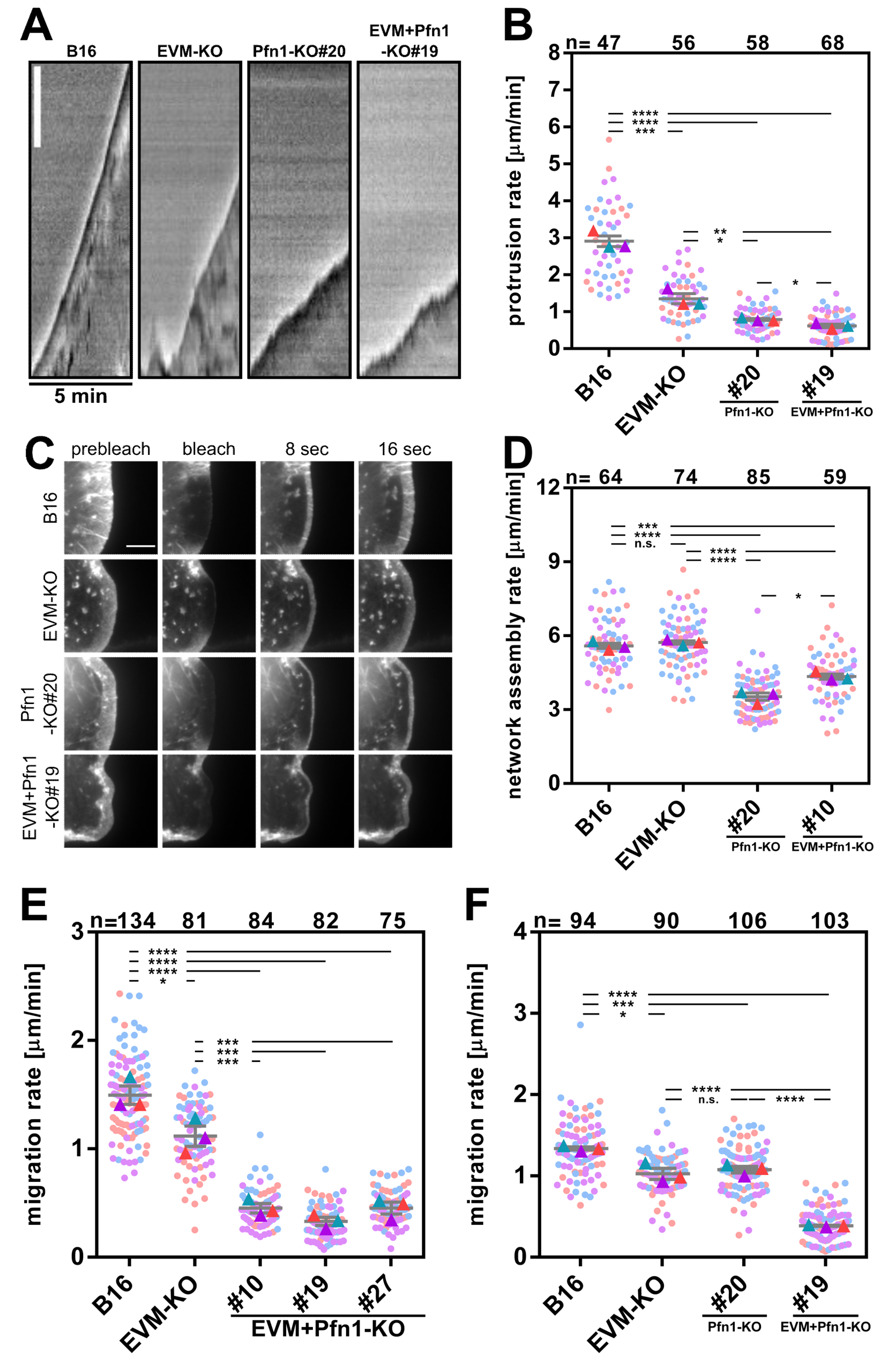
Combined Ena/VASP and profilin removal differentially affects lamellipodial protrusion, actin network assembly and overall random cell migration. (**A**) Side-by-side kymography of representative lamellipodial edges of EVM-KO (clone #23.7.66), Pfn1-KO (clone #20) and EVM+Pfn1 quadruple-KO (clone #19) from data acquired by time-lapse phase contrast microscopy; time is as provided and white scale bar corresponds to 3µm. (**B**) Quantitation of lamellipodial protrusion rates determined from kymographs of individual lamellipodia (shown in A) for the different cell types. The protrusion rates determined for each cell line in three independent experiments are shown as superplots, displaying measurements from individual cells and their arithmetic means as color-coded dots and triangles, respectively. Grey lines and error bars display means and standard errors of the means (SEMs), n = numbers of measured cells. All datasets were statistically compared with each other in all combinations using two sample two-sided T-tests; *p≤0.05; **p≤0.01; ***p≤0.005; ****p≤0.001. (**C**) Fluorescence recovery after photobleaching (FRAP) analysis in lamellipodia of cells with indicated genotype transiently expressing EGFP-tagged actin. As before, fluorescence recovery from the front following the bleach is taken as direct readout of lamellipodium network assembly rate. Scale bar, 5 µm. (**D**) Quantitation of network assembly rate assessed in cell lines as representatively shown in C. Data are displayed as superplots analogous to what’s reported for protrusion rates in B. Statistics: two sample two-sided T-tests; n.s. not significant; *p≤0.05; ***p≤0.005; ****p≤0.001. (**E**) Quantitation of random migration rates of B16-F1 wildtype (B16) *versus* EVM-KO (clone #23.7.66) *versus* the three selected EVM+Pfn1 quadruple-KO cell lines, as indicated. Migration rates were extracted from tracks (see below) over time periods of 10 hours. Data are displayed as superplot data as above and from three independent experiments (distinct colors). Statistics: two sample two-sided T-tests; *p≤0.05; ***p≤0.005; ****p≤0.001. (**F**) Side-by-side comparison of the impact on cell migration of EVM- *versus* Pfn1-removal or both combined using individual cell lines, as indicated. Data are displayed as described for E. For representative movies serving as raw data for the quantitation, see Supplementary Video 8. Statistics: two sample two-sided T-tests; n.s. not significant; *p≤0.05; ***p≤0.005; ****p≤0.001.

**Extended Figure 8.**
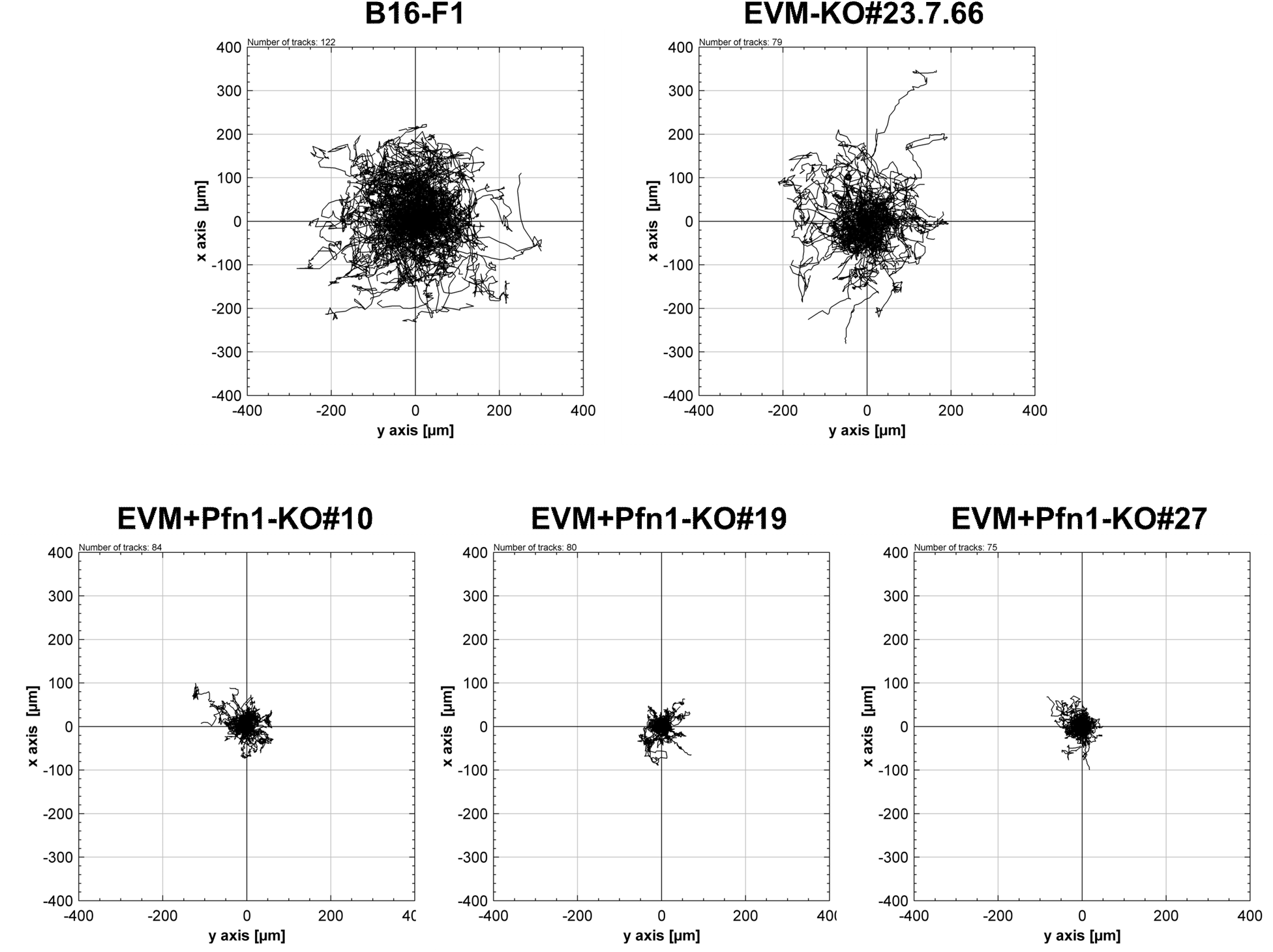
Profilin removal virtually abolishes cell migration in the absence of Ena/VASP family members. Random migration performance of B16-F1 cells lacking EVM alone *versus* both EVM and Pfn1, as indicated. Trajectory plots as obtained from random migration movies of distinct cell lines migrating on laminin. Black trajectories are individually tracked cells, the starting points of which were positioned in the center of the coordinate system. Cells were tracked for a maximum of 10 hours. Note the comparably moderate size reduction of trajectory plots as compared to wildtype in the cell line lacking EVM, consistent with reduced migration rates independently measured in Extended Figs. 7E and –F, but the dramatic compression of trajectory plot sizes in all clones (#10, #19, #27) lacking both EVM and Pfn1.

**Extended Figure 9.**
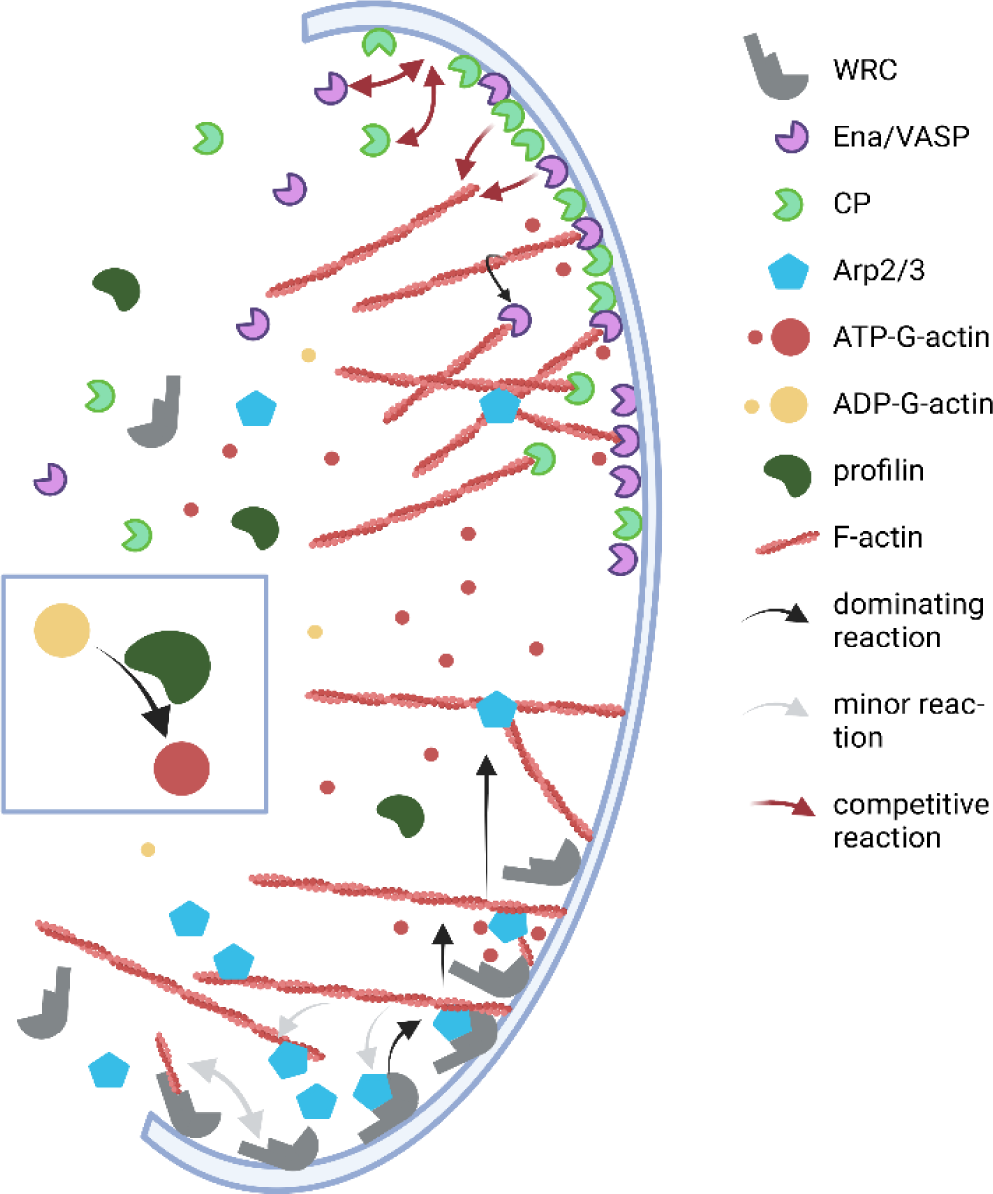
Description of the mathematical model including seven model Figures (Fig. M1-7) and four Tables (Table 1-4).

### Supplementary Video legends

**Supplementary Video 1. Potent filopodia induction by active mDia2 requires profilin.** Time-lapse phase contrast (phase) and fluorescence microscopy, as indicated, of B16-F1 cells transiently transfected with EGFP-tagged, full length mDia2 (EGFP-mDia2-FL) or a constitutively active variant of the formin (EGFP-mDia2-ΔDAD) known to potently stimulate filopodia formation (see e.g. PMID 18755006). As opposed to full length mDia2, which remains inactive and cytosolic in both the presence and absence of Pfn1, EGFP-mDia2-ΔDAD induces dozens if not hundreds of filopodia or filopodia-like structures in the presence, but not absence of Pfn1. Interestingly, cells lacking Pfn1 are capable of prominent mDia2-ΔDAD positioning at the plasma membrane, but emanation of filopodial bundles appears abrogated in these conditions.

**Supplementary Video 2. Genetic disruption of profilin 1 reduces random cell migration.** Time-lapse phase contrast microscopy of B16-F1 wildtype (left) compared to three independently generated Pfn1-KO cell lines, as indicated. Movie was recorded over a time period of 10 hours.

**Supplementary Video 3. Pfn1 gene disruption does not eliminate lamellipodia protrusion.** High magnification time-lapse phase contrast microscopy of representative cells in the presence and absence of profilin 1, revealing protrusion of smooth, convex lamellipodia in cells of both genotypes. Note that Pfn1-deficient lamellipodia appear to protrude more slowly as compared to wildtype cell lamellipodia and to be largely devoid of embedded microspike bundles.

**Supplementary Video 4. Pfn1-deficient lamellipodia are capable of actin filament dynamics turnover.** High magnification time-lapse phase contrast and fluorescence microscopy of representative cells harboring (left panel) or lacking Pfn1, as indicated, and transiently transfected with EGFP-actin followed by seeding on laminin. Note the protrusion of smooth lamellipodial edges in Pfn1-deficient cells lines, which consistently appeared largely devoid of F-actin-rich microspike bundles.

**Supplementary Video 5. Pfn1-deficient lamellipodia display reduced, but not abrogated actin network assembly.** Video microscopy of FRAP of EGFP-tagged actin transiently expressed in wildtype (B16-F1) versus Pfn1-deficient clones (KOs#6, #10, #20). Note the somewhat reduced actin network assembly rate in cell lines lacking Pfn1 as compared to control (left panel).

**Supplementary Video 6. Lamellipodia can incorporate low levels of Arp2/3 complex in the virtual absence of profilin.** Pfn1-deficient cells (clone #20) and WT controls (left) were transiently transfected with EGFP-tagged ArpC5B, one of two variants of the smallest subunits of Arp2/3 complex, formerly known as p16.

**Supplementary Video 7. Genetic removal of Arp2/3 complex in the virtual absence of profilin abrogates random cell migration in B16-F1 cells.** Arp2/3 complex-deficient cells have previously been found capable, in principle, of migration in B16-F1 melanoma and other mesenchymal cell models (Dimchev *et al*., 2021; Suraneni *et al*., 2012; Wu *et al*., 2012), albeit with reduced efficiency as compared to WT. In the absence of Pfn1 and thus the virtually complete elimination of acceleration of formin-dependent actin filament elongation, additional removal of Arp2 was detrimental to the efficiency of cell migration in the two Pfn1+Arp2-KO cell lines analyzed (see right panels). This may well be explained by the removal of lamellipodia due to the absence of Arp2/3 complex activity, but be enhanced in the extent of the phenotype due to the combined elimination of both proper formin and Arp2/3 complex activities.

**Supplementary Video 8. Ena/VASP and profilin contribute to the efficiency of random cell migration in a highly non-redundant fashion.** Comparative time-lapse phase contrast microscopy of distinct cell lines, as indicated. Ena/VASP family-and Pfn1-KO cell lines display at best moderate effects on the efficiency of random cell migration, but the combination of these knockouts (EVM+Pfn1-KO) causes severe defects in migration, comparable perhaps in extent to the combined elimination of Pfn1 with Arp2 (see above).

## The mathematical model

**Figure M1:** Reactions defining the model. WRC translocates from the cytosol to the membrane. It binds Arp2/3 complex and the barbed ends of actin filaments. The WRC-Arp2/3 complex binds to the side of filaments. The WRC-Arp2/3-filament complex forms a branch, the WRC-Arp2/3 bond dissociates without branch formation or the filament dissociates. VASP and capping protein (CP) bind to the membrane and compete with each other for binding sites on the membrane. Membrane-bound CP and membrane-bound VASP also compete with each other for filament barbed ends. CP bound to barbed ends quickly disappears from the tip membrane due to transport with retrograde flow. The VASP- filament complex can dissociate, stay, elongate the filament or be transported away with retrograde flow if elongation is too slow or stopped by some unspecified reason.

The model comprises the dynamic interactions of F-actin filaments and/or their barbed ends with WAVE regulatory complex (WRC), Arp2/3 complex, capping protein (CP), Ena/VASP family proteins (VASP) and the protrusion tip membrane. Fig. M1 illustrates the reactions defining the model. If we include a binding reaction, we also include the corresponding dissociation unless retrograde flow transports the bound molecule away. This applies to CP bound to filament tips and Arp2/3 bound to filament sides.

**Table 1:** Variables of the model.

| Table 1 Variables of the model |  |
| --- | --- |
| variable | description |
| $w_m$ | WRC |
| $wa$ | WRC-Arp2/3 complex |
| $wf$ | WRC-filament complex |
| $waf$ | WRC-Arp2/3-filament complex |
| $f$ | free barbed end |
| $vf$ | VASP bound to barbed ends |
| $vm$ | VASP bound to tip membrane |
| $cm$ | CP bound to tip membrane |

The variable wm is bare WRC bound to the tip membrane. WRC arrives at the tip membrane by a diffusion process described by the first term of the wm-dynamics below with the rate constant s_10_. W is the concentration of wm on the tip membrane that is in equilibrium with the bulk concentration of WRC. The two following terms describe binding (s_1_) of cytosolic Arp2/3 and spontaneous dissociation (s_2_) of Arp2/3 complex. A_bulk_ denotes the (cytosolic) pool of unbound Arp2/3 complexes. Terms 4 and 5 describe the free filament end binding (s_3_) to WRC and its dissociation (s_4_):

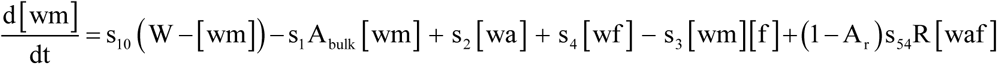

The last term (s_54_) arises from the dynamics of the complex WRC-Arp2/3 bound to a filament (waf). Bonds within this complex are under force due to network assembly and retrograde flow. The faster the network assembly rate R, the sooner the tension on these bonds will rise. Therefore, we set the rate, with which bonds within this complex break, proportional to R. Simulations including a more detailed force balance and an exponential dependency on R provided results very similar to those with a linear R-dependence. On this basis, we chose the simpler linear rate expression. If a bond of the complex WRC-Arp2/3 bound to a filament breaks, the fraction A_r_ breaks between filament and WRC-Arp2/3 and 1-A_r_ between WRC and Arp2/3. If none of the bonds break, a new filament is nucleated and is initially bound to WRC (variable wf). This happens with the rate proportional to s_5_ in the wf- dynamics

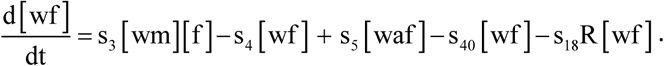

WRC-bound filaments also arise from free ends binding to wm. Here and in all following equations, s_40_ is the rate constant of loss of filaments e.g. due to severing. The term with the rate s_18_R is proportional to the network assembly rate R and describes the loss of WRC-bound filaments from the tip membrane due to transport with retrograde flow. The model fit introduced below showed that both severing and transport contribute little to the dynamics of filament tips at the protrusion tip.

The dynamics of Arp2/3 complex-bound WRC, variable wa, comprises binding of free Arp2/3 (A_bulk_) and its dissociation. The WRC-Arp2/3 complex may bind (s_51_) to the sides of filaments with free ends f, WRC-bound filaments wf, VASP bound filaments vf or filaments from other populations f_e_, like e.g. formin-bound ones (3. term). The dissociation of Arp2/3 complex-filament bonds enters by the last term:

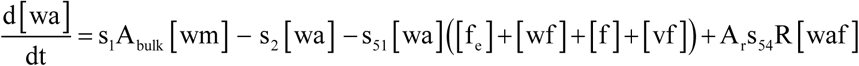

The binding of the WRC-Arp2/3 complex (wa) to a filament entails either formation of a branch (s_5_) or breakage of the WRC-Arp2/3-filament complex (s_54_R):

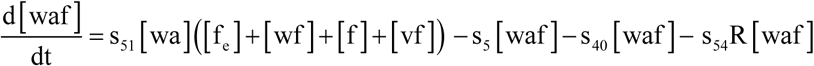

The binding term and the break-term do not show up in the dynamics for filaments, since Arp2/3 complex binds to the sides of filaments belonging to the populations [f], [wf], [vf] [or f_e_], without changing filament numbers in these populations. However, the term branch generation creates a new filament and thus shows up in the [wf]-dynamics.

The first and second terms in the dynamics of free barbed ends [f] describe dissociation from WRC and binding of free barbed ends to WRC. The next two terms represent the same for binding of VASP bound to the tip membrane (s_70_, s_71_). The term with the rate s_82_ describes capping by CP bound to the tip membrane cm. Dissociation of CP from filament tips does not contribute to the [f] dynamics, as we assume that capped barbed ends are immediately transported away with retrograde flow. The term with the rate s_38_R describes the loss of free filament ends from the tip membrane due to advection with retrograde flow:

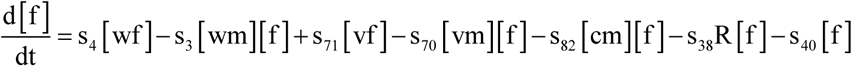

The experimental results comprised the interesting observation of an increase in leading edge VASP accumulation upon Pfn1 knockout, in spite of decreasing F-actin density. This implies the existence of VASP binding sites at the membrane (m) in addition to barbed ends. We denote those, occupied VASP-binding sites vm. A competition between VASP and CP at the leading edge of lamellipodia was previously proposed in the context of competition for barbed end binding. This competition is experimentally supported by the observation of increased CP accumulation in the absence of Ena/VASP (Damiano-Guercio *et al*, 2020), and the *vice versa* experiment (this study, Extended Fig. 6E, F, G). This means that VASP intensity at the lamellipodium tip goes down when CP goes up and *vice versa*. The existence of VASP binding sites at the tip membrane renders such a competition impossible unless VASP and CP compete not only for barbed ends, but also for these tip membrane binding sites m. Denoting the binding sites occupied with CP cm, the following dynamics realize such a competition:

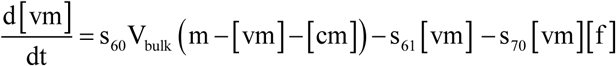

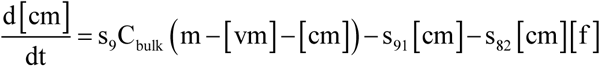

V_bulk_ and C_bulk_ denote the cytoplasmic concentrations of VASP and CP, respectively. We have chosen here the simplest possible dynamics recruiting VASP and CP to the membrane, since we lack detailed information on the process. Those dynamics are able to reproduce the experimental results in good approximation. However, the recruitment might be more complicated and might comprise additional steps. The term with s_61_ describes VASP dissociation back into the cytosol. The experimental data show an increase of VASP at the lamellipodium edge concomitant with a decrease of F-actin density (see above). We thus suggest that VASP binding to barbed ends causes a flux of VASP out of the vm- population. We assume on this basis that VASP dissociating from barbed ends does not return to the membrane binding sites, i.e. to vm, immediately, but instead diffuses back into the cytosol. The last term describes binding of membrane-bound VASP to barbed ends.

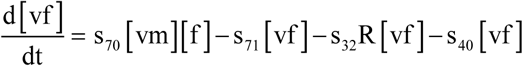

The dynamics of barbed ends bound to membrane-bound VASP also includes a transport term. The other terms in the vf-dynamics are free end binding and dissociation.
Experimental results are provided as concentration ratios relative to wildtype. We lack information on absolute concentrations. Consequently, we scale all dynamic model variables with W. This scaling does not affect concentration ratios. We are exclusively considering stationary states. Therefore, we cannot make any statements on absolute rate values and consequently scale time with s_10_. Thus, we obtain dimensionless parameter values by the scaling:

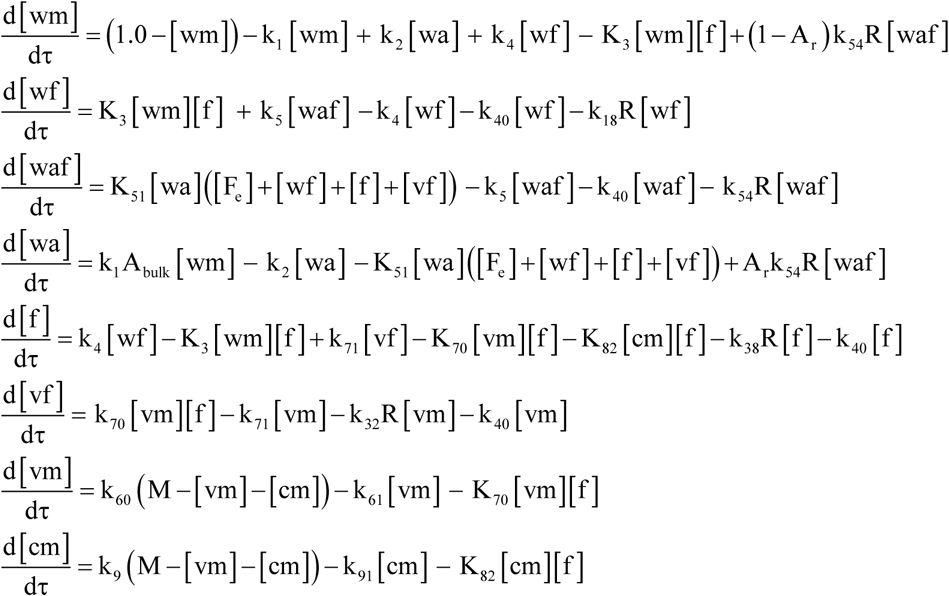

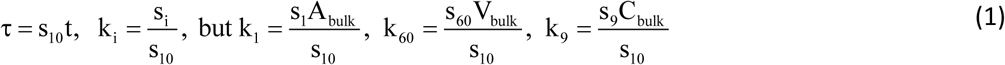

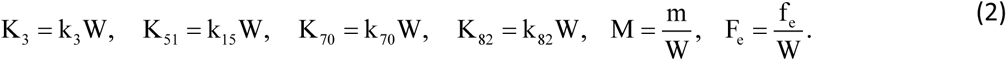

Here, the names for the scaled dynamic variables have remained like the unscaled ones and the parameter names have been changed only for those affected by the scaling. The fit to the experimental results has been done with the scaled model, which is possible since all results are concentration ratios.

**Table 2:** Parameters of the model.

| Table 2: Parameters of the model |  |  |
| --- | --- | --- |
| parameter | description | scaled parameter |
| $s_1$ | binding rate of Arp2/3 to WRC | $k_1 = s_1 A_{\text{bulk}}/s_{10}$ |
| $A_{\text{bulk}}$ | Arp2/3 cytosolic concentration | |
| $s_{10}$ | equilibration rate of total WRC on the membrane | |
| $s_{18}$ | rate of WRC-filament complex loss due to retrograde flow | $k_{18} = s_{18}R/s_{10}$ |
| $s_2$ | dissociation rate of Arp2/3 complex from WRC | $k_2 = s_2/s_{10}$ |
| $s_3$ | binding rate of filaments to WRC | $K_3 = Ws_3/s_{10}$ |
| $Rs_{32}$ | rate of VASP-bound barbed end loss due to retrograde flow | |
| $Rs_{38}$ | rate of barbed end loss due to retrograde flow | |
| $R$ | network assembly rate | |
| $s_4$ | dissociation rate of filaments from WRC | $k_4 = s_4/s_{10}$ |
| $s_{40}$ | rate of filament loss due to severing, bundling, and unspecified processes | $k_{40} = s_{40}/s_{10}$ |
| $s_5$ | rate of branch formation | $k_5 = s_5/s_{10}$ |
| $s_{51}$ | binding rate of Arp2/3-WRC complexes to filaments | $K_{51} = Ws_{51}/s_{10}$ |
| $Rs_{54}$ | dissociation rate of WRC-Arp2/3-filament complexes without branch formation | $k_{54} = s_{54}/s_{10}$ |
| $(1-A_r) Rs_{54}$ | dissociation rate of Arp2/3-filament complexes and WRC without branch formation | |
| $A_r Rs_{54}$ | dissociation rate of filaments and Arp2/3-WRC complexes without branch formation | |
| $A_r$ | fraction of dissociation events of WRC-Arp2/3-filament complexes without branch formation occurring at the Arp2/3-filament bond | |
| $s_{60}$ | rate of VASP binding to tip membrane | $k_{60} = s_{60} V_{\text{bulk}}/s_{10}$ |
| $s_{61}$ | rate of VASP dissociation from tip membrane | $k_{61} = s_{61}/s_{10}$ |
| $s_{70}$ | rate of membrane-bound VASP binding to barbed ends | $K_{70} = Ws_{70}/s_{10}$ |
| $s_{71}$ | rate of VASP dissociation from barbed ends | $k_{71} = s_{71}/s_{10}$ |
| $s_{82}$ | rate of membrane-bound CP binding to barbed ends | $K_{82} = Ws_{82}/s_{10}$ |
| $s_9$ | rate of CP binding to tip membrane | $k_9 = s_9 C_{\text{bulk}}/s_{10}$ |
| $s_{91}$ | rate of CP dissociation from tip membrane | $k_{91} = s_{91}/s_{10}$ |
| $f_e$ | filaments bound to formins or other membrane proteins | $F_e = f_e/W$ |
| $V_{\text{bulk}}$ | cytosolic concentration VASP | |
| $C_{\text{bulk}}$ | cytosolic concentration of CP | |
| $m$ | concentration of binding sites for VASP and CP on the tip membrane | $M = m/W$ |
| $W$ | equilibrium concentration of WRC on the tip membrane | |

## Fitting experimental results

The experimental results constitute a set of measured ratios of densities of lamellipodial factors in various conditions close to or at protrusion tip membranes. Molecules which are being transported away with retrograde flow from the tip membrane contribute to the fluorescence intensity as long as they are close to the tip membrane. Therefore, we include them in the ratio calculation. The density of these molecules in the F-actin network is given by the rate with which they bind to the network divided by the network assembly rate R. Filament tips transported with retrograde flow contribute on average one half of their density to the F-actin density, since the filament does not contribute between barbed end and protrusion tip membrane to F-actin density. Thus, we obtain for total amounts

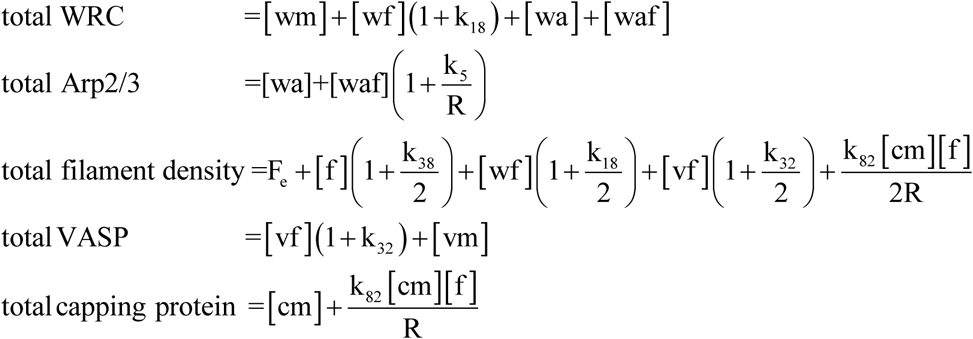

We model Pfn1-KO by reducing the branching rate k_5_ by the factor f_pfn_. The branching rate is proportional to the square of the concentration of polymerizable G actin according to Carlsson and colleagues (Carlsson, 2005; Carlsson *et al*, 2004). Hence, the factor by which polymerizable G-actin is reduced in Pfn1-KO is 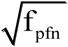. We model Ena/VASP- and CP-KO by setting the values of V_bulk_ and C_bulk_, respectively, to 1% of their control values. Consequently, k_60_ and k_9_ have 1% of their control values in those knockouts. Setting them to 0 would have rendered the Jacobi matrix of the system singular, which we chose to avoid since we also used implicit integration methods and Newton Raphson methods to determine stationary states (see section Numerical methods). The R-value has been experimentally determined.

## Results

Table 3 displays 13 values to which we can fit the mathematical model. Additionally, we know that VASP increased 2-3fold upon CP KO, such that 14 values need to be fit by the model outcome. The network assembly rate is a parameter in our modelling (Table 4).

**Table 3:** Experimental results constituting fit criteria for the model.

| Table 3: Experimental results constituting fit criteria for the model |  |  |  |
| --- | --- | --- | --- |
|  | EVM-KO ratio with control | Pfn1-KO ratio with control | EVM+Pfn1-KO ratio with control |
| F-actin | 0.506 | 0.495 | 0.303 |
| Arp2/3 | 1.29 | 0.491 | 0.397 |
| WRC | 0.934 | 0.677 | 0.500 |
| CP | 1.71 | 0.602 | 1.80 |
| VASP/Mena | 0.0 | 3.20 | 0.0 |

**Table 4.** Experimental results for the network assembly rate (parameter R)

|  |  |  |  |  |
| --- | --- | --- | --- | --- |
|  | control | EVM-KO | Pfn1-KO | EVM+Pfn1-KO |
| network assembly rate | 6.48 $\mu\text{m min}^{-1}$ | 5.3 $\mu\text{m min}^{-1}$ | 3.18 $\mu\text{m min}^{-1}$ | 3.81 $\mu\text{m min}^{-1}$ |

**Figure M2.**
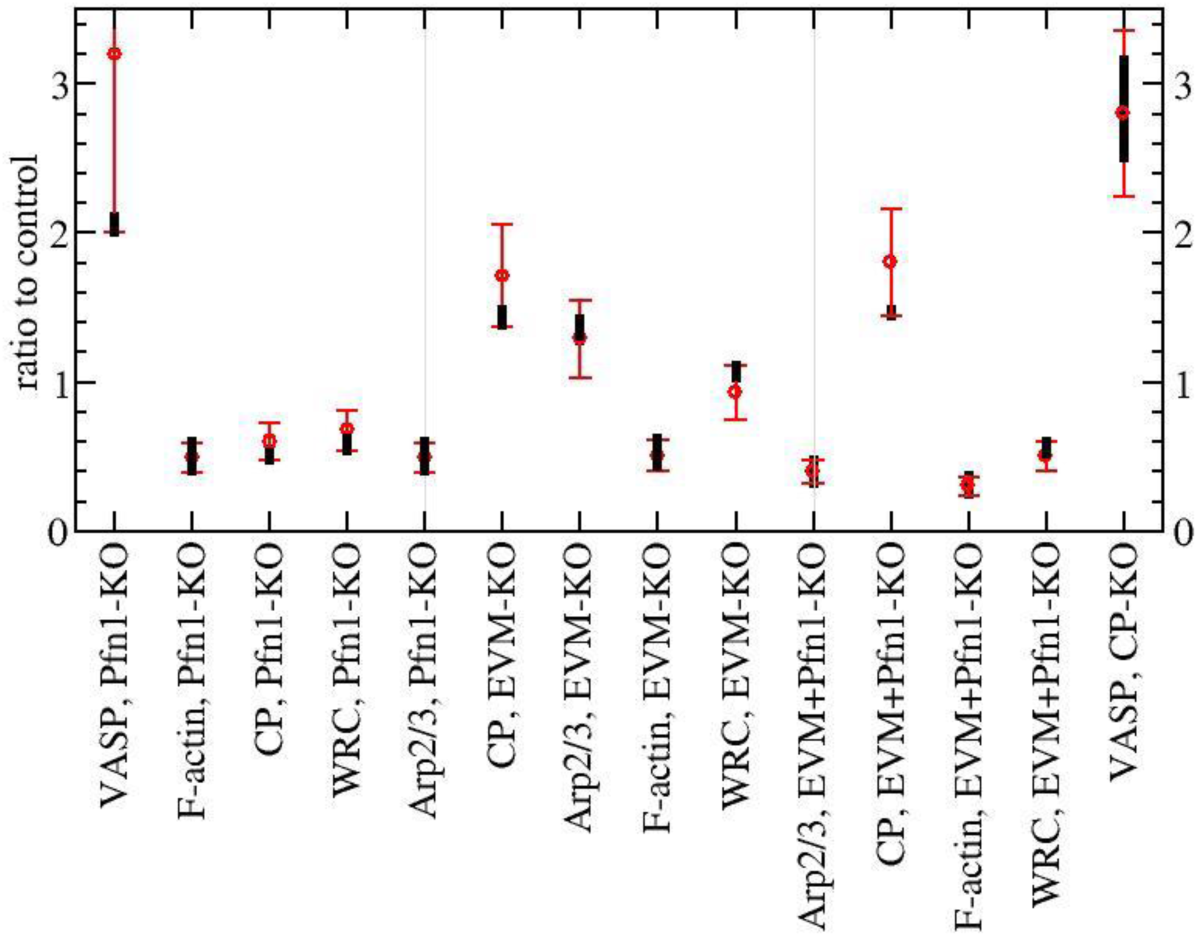
Fit results for the 14 fit criteria. We considered a criterion as met if the model output was in the range indicated by the respective, red line. The model output ranges represented by black lines in each case arose from 2218 parameter value sets each meeting the criteria. They were found in a sample of 5 10^9^ parameter value sets.

We defined a fit criterion as met if the corresponding model variable output was within +/-20% of each value listed in Table 3. VASP increased 3.2fold and Mena 1.67 upon Pfn1-KO. Within this range, we defined the criterion (called VASP Pfn1-KO in Fig. M2) met if model output was larger than 2.0. We find values up to about 2.2 as simulation output (but see next section also). We found parameter value sets producing outputs within our 20%-range for all the other criteria.

**Figure M3.**
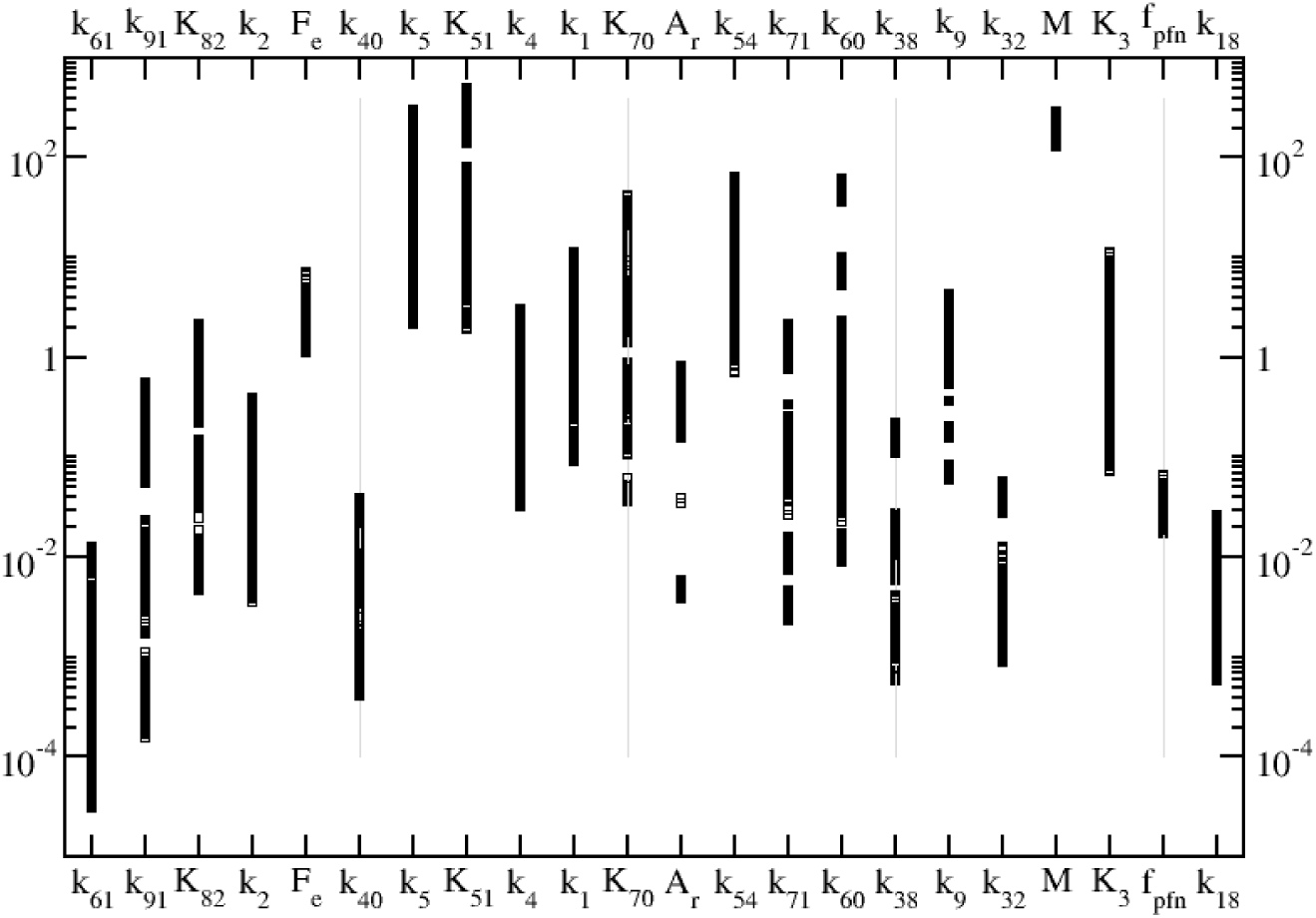
Values of the scaled parameters defined in Eqs. 1,2 resulting from the fits in Fig. M2. The model output ranges represented comprise 2218 parameter value sets each meeting the criteria. They were found in a sample of 5 10^9^ parameter value sets.

**Figure M4.**
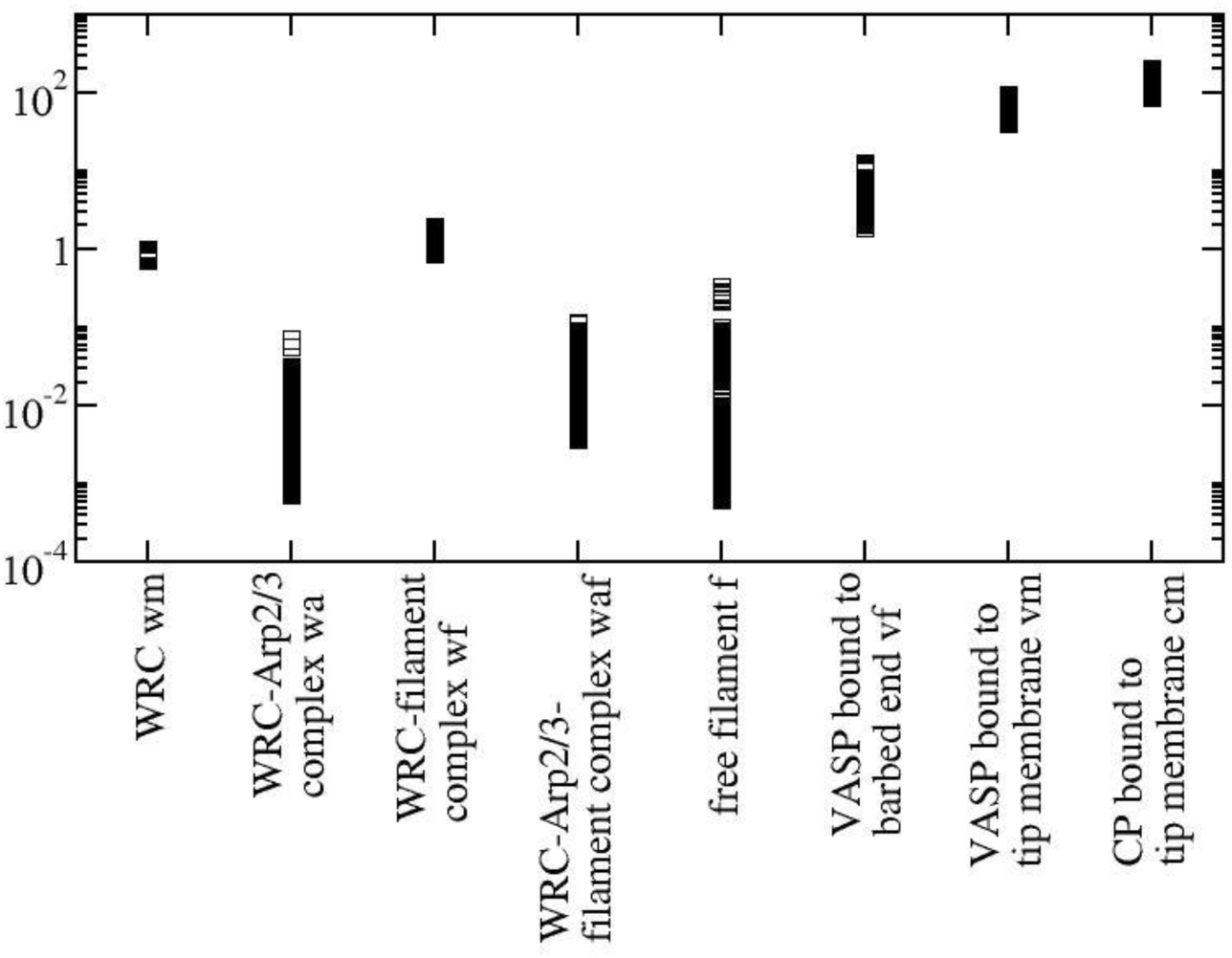
The values of the scaled control variables resulting from the model fits in Fig. M2. The model output range represented comprises 2218 parameter value sets each meeting the criteria. They were found in a sample of 5 10^9^ parameter value sets.

The parameter values and variable values resulting from the fits are shown in Figs. M3 and M4. We find fits to the fit criteria in Fig. M2 for rather large ranges of parameter values. Hence, the mechanism formulated in the model does not depend on very specific assumptions on parameter values (see parameter value ranges in Fig. M3), but instead is rather robust.

The largest fraction of filaments is VASP-bound, followed by WRC-bound. Filaments with WRC-bound Arp2/3 bound to their sides are in the range below 1% of total filament numbers. Filaments with free barbed ends in the range below 2%.

The fastest scaled rate in Fig. M3 is K_51_, the binding of WRC-bound Arp2/3 to filament. The parameter f_pfn_ describes the modelling of Pfn1-KO. We model it as a reduction of the branching rate s_5_ by this factor. Its value is in the range of 0.017-0.083. The branching rate is proportional to the square of the concentration of polymerizable G actin (Carlsson, 2005; Carlsson *et al*., 2004). Hence, the factor by which polymerizable G-actin is reduced by Pfn1 KO is 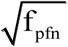 corresponding to a reduction of polymerizable G-actin to about 13.0%-28.7% of its control value in Pfn1-KO. The expression of actin in Pfn1-KO cells was measured to be reduced to between 35% and 42%, dependent on the Pfn1-KO clone (main text Fig. 1G). Hence, mean percentages are a factor 1.22-1.46 higher than the upper limit of the range suggested by the model, but some individual experimental values are in the range of model predictions (Fig. 1G).

The slowest rate is the spontaneous dissociation rate k_61_ of VASP from the membrane binding site m. The spontaneous dissociation of CP from the membrane binding site is also slow. Binding of VASP and CP with the rates k_60_ and k_9_ resp. to m is also slow. The binding of VASP and CP to barbed ends has faster rates (K_70_ and K_82_) than binding of VASP and CP to m. Therefore, most of the membrane-bound VASP and CP molecules will bind barbed ends, at least with control values of barbed end density.

Other slow processes, i.e. processes not dominating the dynamics, are the losses due to transport by retrograde flow of WRC bound to filament (k_18_), VASP bound to filament (k_32_), free barbed ends (k_38_) and filament loss due to severing (k_40_).

The parameter A_r_ describes the fraction of bonds in the WRC-Arp2/3-filament complex breaking between WRC and Arp2/3 upon stress. In principle, it can assume values between 0 (no dissociation of the filament from the WRC-Arp2/3-filament complex) and 1 (no dissociation of the WRC from the WRC- Arp2/3-filament complex). Fig. M3 shows that we find fits for the whole range of values. Hence, the mechanism formulated in the model does not depend on assumptions on this fraction.

## Lower margin for fit criterion on ratio of CP upon Pfn1-KO to control

If we allow for smaller values of the ratio of CP upon Pfn1-KO to control than our 20% criterion, we find larger values of the ratio of VASP upon Pfn1-KO to control, i.e. up to 2.62 (Fig. M5). This confirms the anti-correlation between VASP and CP. Parameter values and variable values resulting from the fit criteria in Fig. M5 are shown in Figs. M6 and M7. They cover slightly larger ranges than in Figs. M3 and M4.

**Figure M5.**
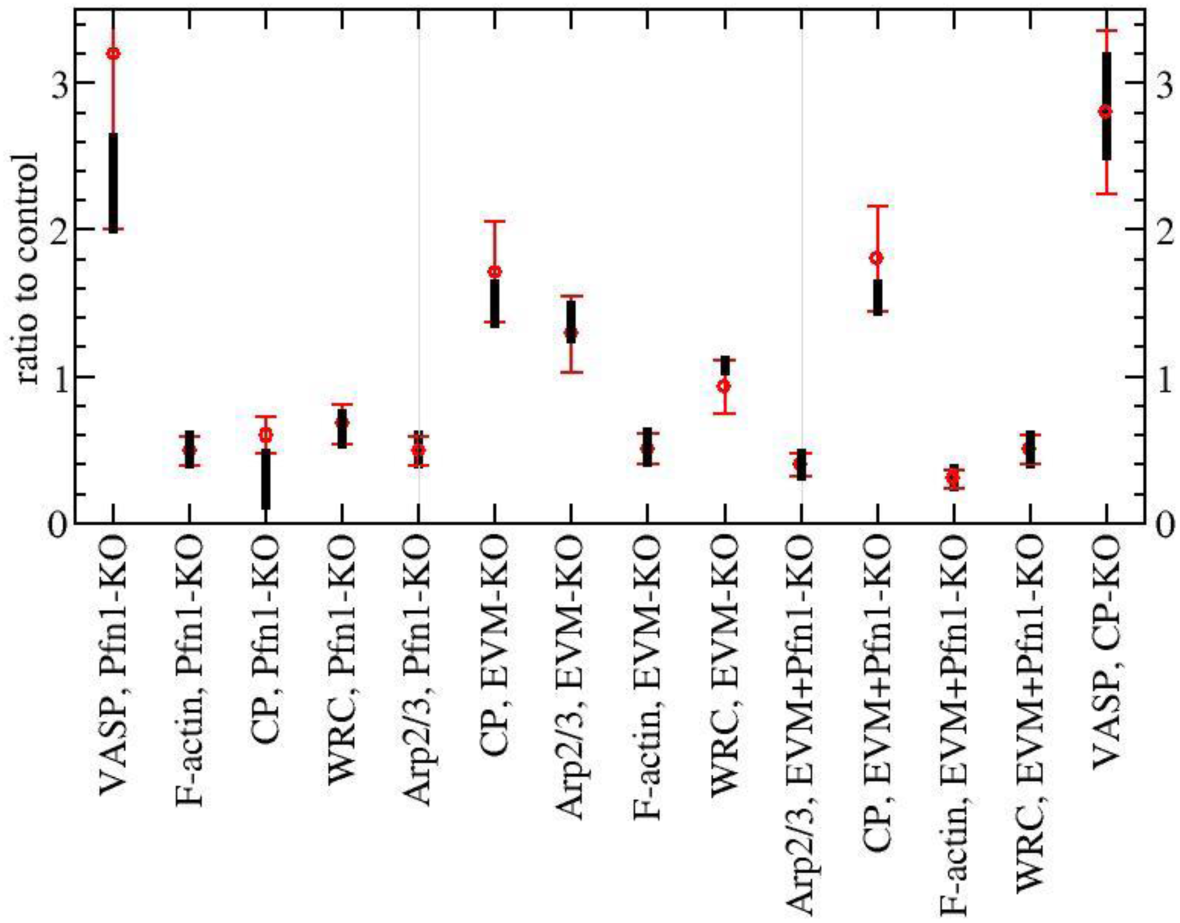
Fit results for the 14 fit criteria. We considered a criterion as met if the model output was in the range indicated by the red lines, except for the criterion ‘CP, Pfn1 KO’. We show here results with this criterion being below the acceptance range. The model output range represented by black lines comprises 169.994 parameter value sets each meeting the criteria. They were found in a sample of 5 10^9^ parameter value sets.

**Figure M6.**
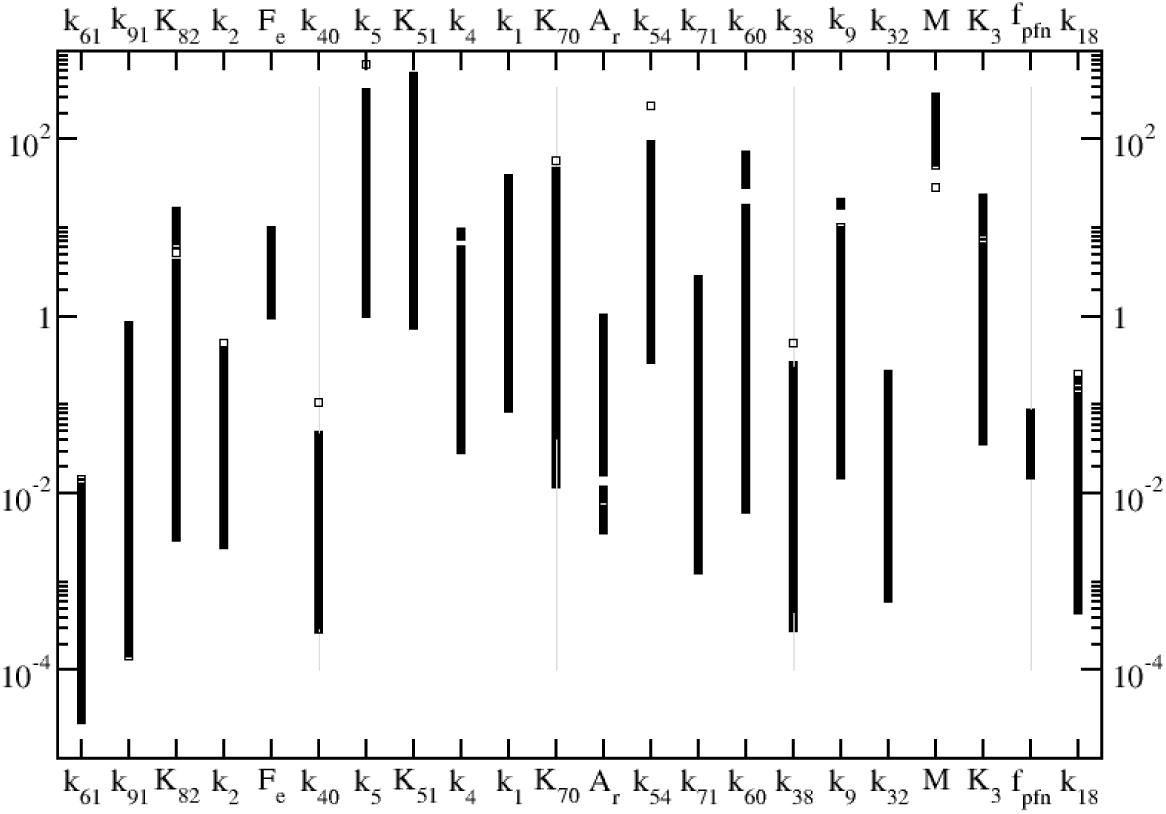
Values of the scaled parameters defined in Eqs. 1,2 resulting from the fits in Fig. M5.

**Figure M7.**
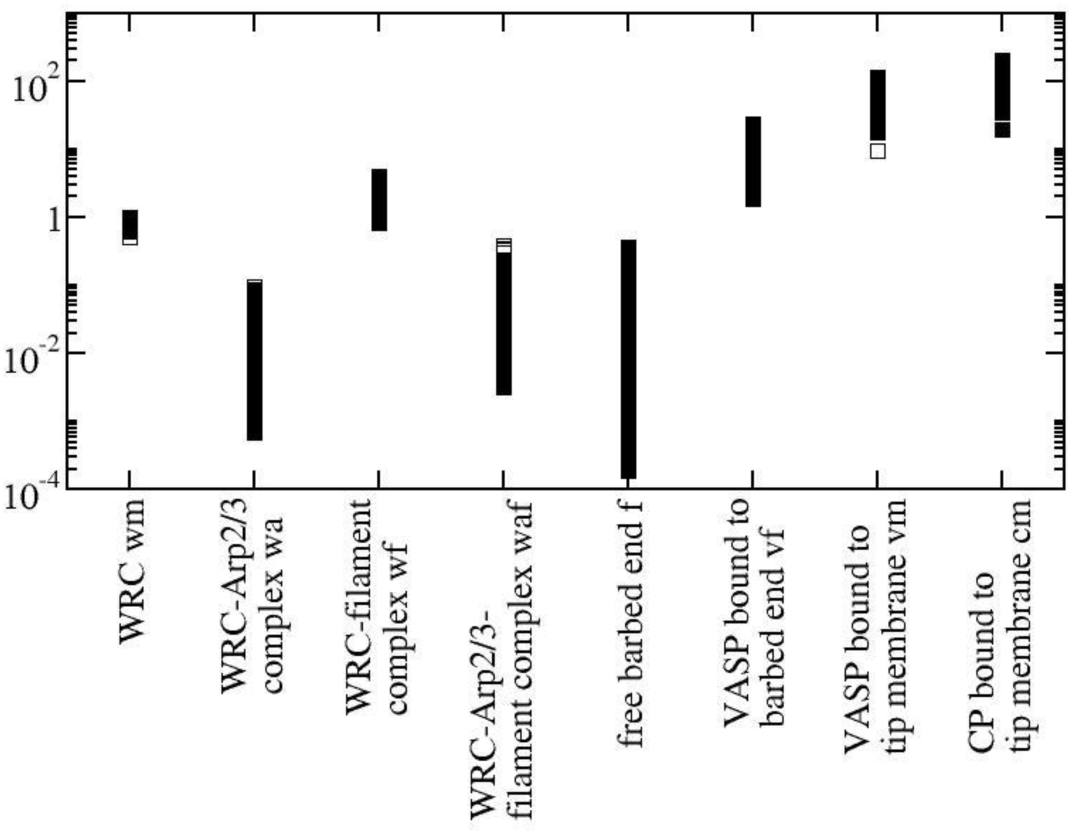
The values of the scaled control variables resulting from the model fits in Fig. M5.

## Numerical methods

Most stationary states were found with a Newton-Raphson iteration (Press *et al*, 2007). The stability of selected stationary points found with this method was checked by integration of the ODEs. If Newton-Raphson iterations did not converge, we alternated integration of the ODEs and Newton-Raphson till convergence was reached. Computations have been carried out on the high performance compute cluster of the Max Delbrück Center for Molecular Medicine Berlin.

